# Gene library deep sequencing for protein super-family profiling

**DOI:** 10.1101/2025.10.20.682913

**Authors:** Rahkesh T Sabapathy, Kara Henry-Cocks, Jackson Feng, Shashikanth Marri, Gustavo Bracho Granado, Harald Janovjak

## Abstract

Unravelling protein function at scale remains challenging. Gain-of-function (GOF) screens harnessing large open reading frame (ORF) libraries enable systematic functional annotations of cell behavior drivers and drug targets. Currently, few sequencing methods are established to analyze existing ORF libraries and facilitate future library generation. Here, we developed a barcode-free deep sequencing method termed pooled plasmid library sequencing (PPLseq). PPLseq achieved high efficiency and single nucleotide accuracy, including the detection of previously undocumented variants, on a prototypical super-family ORF library (>300 human G-protein-coupled receptors (GPCRs)). We next combined PPLseq with high-throughput gene engineering to generate a new library of 246 GPCR-fluorescent protein (FP) fusions. We quantified expression of all members of this library and identified robustly expressed yet understudied receptors. Collectively, we demonstrate accessible, scalable, and sensitive ORF library sequencing towards a deeper understanding of proteome function.

## Introduction

A major scientific endeavor is to understand the form and function of the ∼25’000 proteins encoded in the human genome. Whereas recent deep learning-based advances, such as AlphaFold [1, 2] or RoseTTAFold [3], provide proteome-wide models of form, defining function most commonly relies on experimental inquiry. Complementary to loss-of-function experiments (e.g., gene knock-out or -down using CRISPR or RNAi), GOF screens allow high-throughput analysis of ORF (over)expression in the context of complex cell states and behaviors ([4–7], and below). However, the generation, propagation, and application of pooled or arrayed ORF libraries is challenged by undocumented and/or undesired coding sequence alterations and non-uniform coverage, as well as recombination events and off-target effects during gene delivery [8–13]. These confounders can individually or collectively limit library reliability and screen power and hence call for rigorous sequence characterization.

Limited large-scale sequencing was performed on ORF libraries prior to the broad availability of NGS methods [14–19]. More recently, NGS verification has been applied to libraries of short regulatory sequences or isolated protein domains [4, 20–23] (typically <300 base pairs (bp)) but less so to full-length ORF libraries including those obtained commercially or from repositories [10, 12, 24]. NGS has been demonstrated to be essential to quantify enrichment in elegant pooled screens but in most studies ORF integrity was not assessed [5, 7, 13, 25–27]. Thus, the potential of NGS to systematically characterize ORF libraries is underexplored.

One particular research area that relies on ORF libraries and thus may benefit from improved methods are studies of large protein (super-)families, such as membrane proteins [30–32]. For instance, ∼300 diverse receptors (and ∼400 olfactory receptors) form the GPCR super-family [33, 34]. GPCRs are expressed in virtually every human cell type, sense diverse ligands, and are the target of ∼30% of all prescription drugs [35, 36]. Only very few super-family-wide GPCR libraries are available and large-scale functional studies of GPCRs remain challenging. Inoue *et al.* and Avet *et al.* determined downstream coupling profiles for 148 or 100 therapeutically-relevant wild-type GPCRs [37, 38]. Kroeze *et al.* fused 314 DNA sequence-optimized GPCRs to a cleavable transactivator domain for high throughput functional assays, and Lv *et al.* and Tedman *et al.* generated extensive libraries of 826 or 940 human GPCRs and splice variants and applied these to surface expression analysis [39–41]. Finally, we generated a library multi-fusion chimeric Class A orphan GPCRs for optogenetics [42]. Sequencing needs to be en par in scale and throughput with high-throughput gene engineering methods, such as those employed in these studies. This will enable the generation of further libraries of GPCRs or even larger protein super-families (e.g., kinases or transcription factors) and ultimately dissect commonalities and differences in super-family-omes.

Here, we systematically developed protein super-family library NGS. We first evaluated whether short-read sequencing, being the most economical and large-scale NGS method, is suited to characterize prototypical ORF libraries. We then developed the experimental workflow termed PPLseq and validated it on a previously established GPCR library. We showed that PPLseq can report library coverage and accuracy with single nucleotide resolution, including previously undocumented ORF variants. We expanded on the use of PPLseq to generate a large fluorescently-labelled GPCR library. Analysis of this library using single-cell confocal microscopy quantified expression levels, including of understudied receptors, as targets for future structure-function analysis.

## Results

### Barcode-free ORF library sequencing

We first asked if short-read NGS is suited for ORF library sequencing without barcoding (**Figure 1a, Supplementary Figure S1**). ORF-specific barcoding prior to NGS library preparation assists with read mapping but is incompatible with the analysis of existing pooled libraries and adds additional processing steps (**Supplementary Figure S1**). Barcodes are not required if library genes are sufficiently different for reads to be mapped to the correct ORF (**Figure 1a**). We tested this computationally for three large human protein super-families: the largest receptor family (314 unique non-odorant GPCRs), the largest enzyme family (495 unique kinases), and a diverse super-family (1993 unique transcription factors (TFs)) (for sequences, source materials, and deduplication methods see **Supplementary Table 1**, **Supplementary Data S1-3** and **Materials and Methods**; the combined lengths of the ORFs in these libraries are 0.42 (GPCRs), 0.88 (kinases) and 3.2 Mb (TFs)). To test for unambiguous read mapping, we searched the libraries for non-unique stretches using four sliding windows (50, 75, 150, or 300 bases (b)) which reflect cycle number and read lengths presets in state-of-the-art short-read sequencers. The emerging data tracks report non-uniqueness akin to ‘uniquome’ analysis in whole genome sequencing [43, 44]. We found that sequences in the GPCR library were unique for all window lengths (**Figure 1b,c**). For window lengths of 150 and 300 b, but not shorter lengths, only a small number of non-unique segments were identified both in the kinase and TF libraries (**Figure 1d-g**, **Supplementary Figure S2**). The kinase library contained only two non-unique ORF pairs which correspond to gene splice variants of MAP2K2 and MST4 (**Supplementary Table S2**). These pairs share eight (150 b) or six (300 b) overlapping segments (**Figure 1d,e**; segment lengths are specified in **Supplementary Figure S2**). In the TF library, ten sequence pairs (150 b) or one sequence pair (300 b) were identified containing a total of 20 (150 b) or two (300 b) overlapping segments (**Figure 1f,g**, **Supplementary Figure S2**). These pairs correspond to paralogs (e.g., *STAT5A* and *STAT5B* [45]), genes that share exons (*ZIM2* and *PEG3* [46]), or genes with internal tandem repeats (e.g., *IFI16* [47]) (**Supplementary Table S2**). Thus, with the exception of a small number of closely related ORFs, correct read mapping is likely at read lengths ≥150 b. Under the assumption that any technical replicate clones of the same ORF are placed in different sequencing pools (termed sub-libraries, see below), barcode-free pooled library short-read sequencing is suited for protein super-family sequencing.

**Figure 1.**
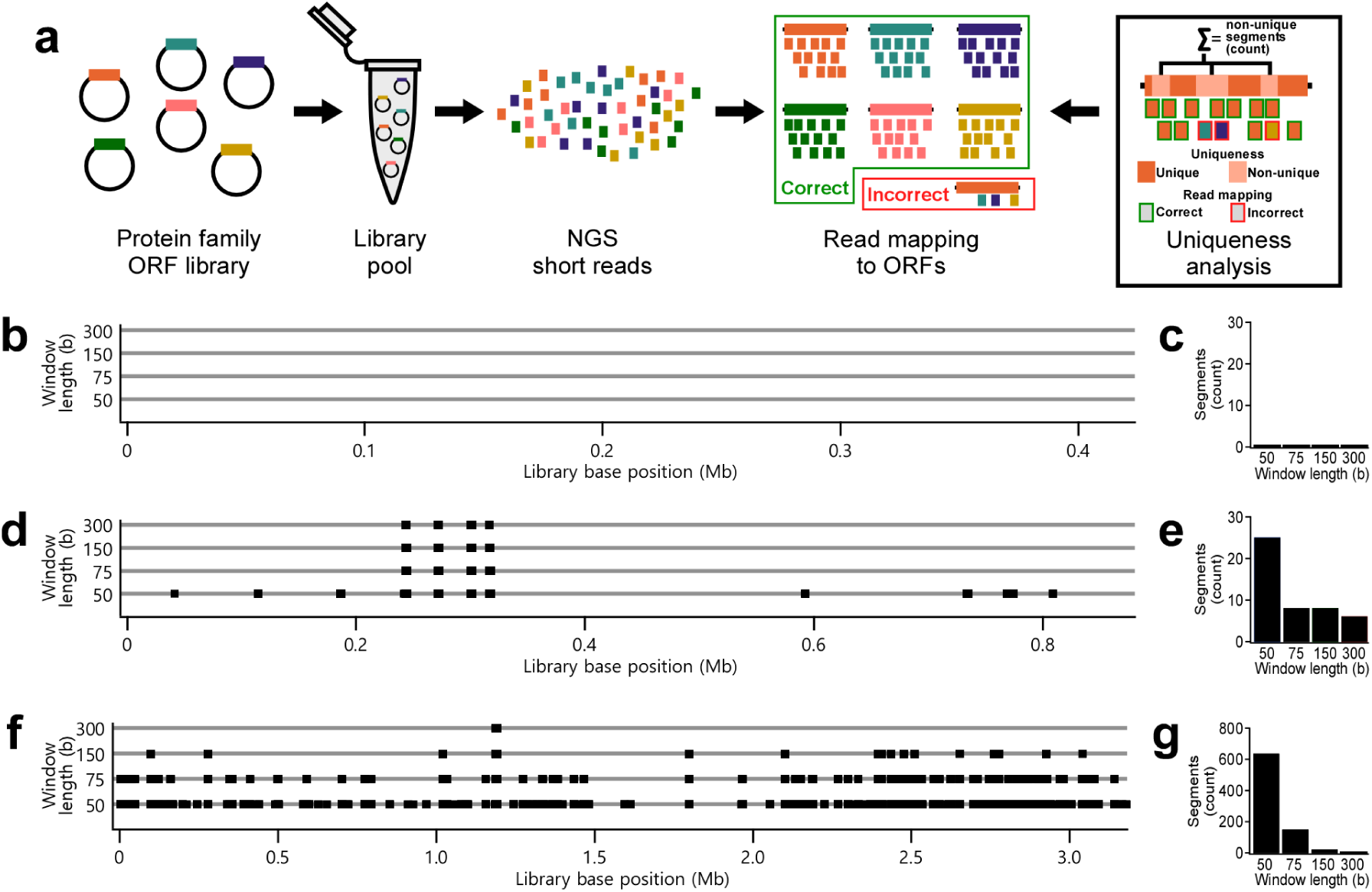
Multi-mapping reads in three large human gene families. (a) Pipeline encompassing plasmid library pool NGS and read mapping to correct (green box) or incorrect (red box) ORFs. Uniqueness analysis identifies non-unique segments that may result in ambiguous read mapping (reads with red outline). (b,c) Uniqueness tracks and the number of non-unique segments in the human GPCR library. (d,e) Uniqueness tracks and the number of non-unique segments in the human kinase library. (f,g) Uniqueness tracks and the number of non-unique segments in the human TF library. Grey horizontal lines in (b,d, and f) represent ORF library sequences from the first base of the first ORF to the last base of the last ORF. Black squares denote non-unique sequence stretches for the window length specified by the Y-axis and at the base position specified by the X-axis. Bars in (c,e, and g) denote the number of continuous non-unique sequence segments identified in the corresponding libraries.

### *PPLseq* workflow

We next established the experimental PPLseq workflow (**Figure 2a**). We focused on the GPCR library because we have reengineered these receptors in the course of this work and because high quality reference sequences are available for 314 unique full length receptors in the PRESTO-Tango library [39]. To generate a library pool, we first replica-plated the cryo-preserved *E. coli* cultures onto agar trays for overnight growth (**Figure 2a**). Growth was observed for all but four genes (**Supplementary Figure S3**, also see below). The localized inoculation spots were pooled and plasmid DNA was isolated using standard silica columns. We performed NGS with a target data yield of 10 to 20 Gb as this would allow for deep sequencing (depth >1000; including plasmid backbones, the library spans ∼2.5 Mb). Sequencing should thus provide complete library deep sequencing with a read excess to compensate for *E. coli* plasmid copy number variability (see below). We chose a WGS library preparation method as it is designed for double-stranded DNA and offers read lengths supported by above uniqueness analysis in paired-end sequencing (150 b). We processed >43 million filtered NGS reads. The main steps of the pipeline were read mapping to ORFs collected in a reference sequence inventory, read statistics (i.e., overall coverage of the sequence from first to last codon or per base sequencing depth), and variant calling (i.e., deviations from the expected sequences) (**Figure 2a**). Pipeline outputs are discussed in the context of ORF recovery and accuracy.

**Figure 2.**
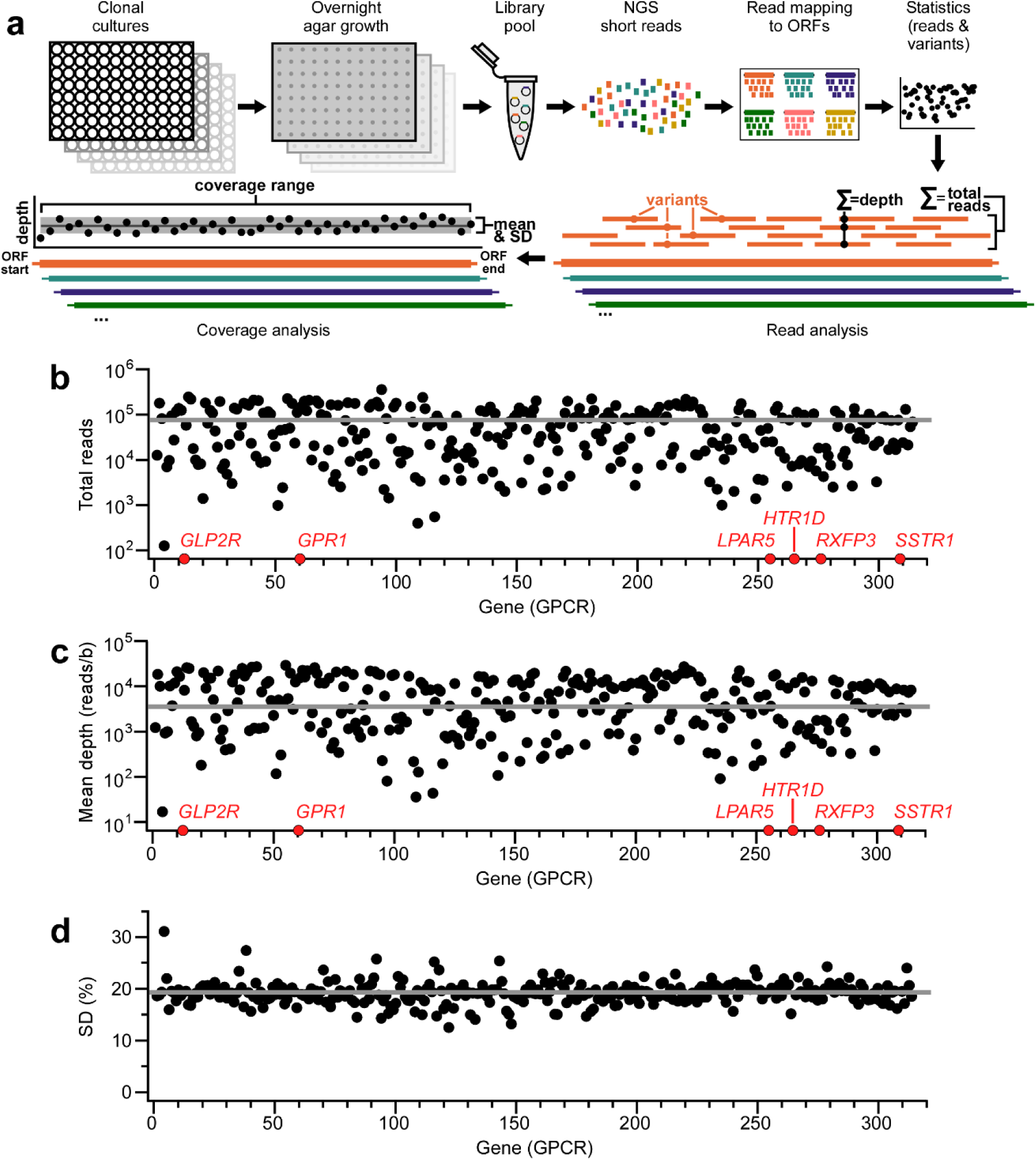
Gene recovery in PPLseq. (a) PPLseq workflow encompassing replica growth of clonal cultures, DNA pooling, NGS, and sequencing statistics. (b) Total reads mapped to each GPCR ORF. Red spheres indicate genes without mapped reads, except for *SSTR1* that was mapped by two reads. (c) Mean depth for each GPCR ORF. Red spheres as in (b). (d) Standard deviation of mean depths of (c). In (b-d), grey lines indicate medians.

### ORF recovery in PPLseq

We assessed recovery in the PPLseq workflow by comparing the identified GPCR ORFs to those expected in the library. We found that reads could be mapped to all but six ORFs indicating an initial gene recovery rate of >98% (**Figure 2b**, **Supplementary Data S4**). These 308 mapped ORFs were sequenced with median counts of 38’487 reads per ORF and with a median depth of 4’284 (**Figure 2b,c**). We noticed that reads and mean depth were not uniformly distributed across the genes (**Figure 2b,c**) or within each gene (**Figure 2d**, **Supplementary Figure S4**), but this did not negatively impact analysis and is discussed below. To assess recovery more completely, we further investigated the six genes with zero or less than two mapped reads (*GLP2R*, *GPR1*, *LPAR5*, *HTR1D*, *RXFP3*, and *SSTR1*; red spheres in **Figure 2b,c**). The library strains that should encode *GLP2R*, *GPR1*, *HTR1D*, and *RXFP3* did not yield inoculation spots in our library cultures and could also not be isolated from the pooled library using polymerase chain reactions (PCR) reactions (**Supplementary Figure S5**). Thus, these genes were likely absent from our preparation of the source library and accordingly not captured by PPLseq. The strains that should encode *LPAR5* and *SSTR1* did yield inoculation spots and plasmids isolated from this growth were analyzed using Sanger sequencing. In the case of the plasmid expected to encode *LPAR5*, Sanger sequencing reported a variant of the human *LPAR5* receptor with an alternative DNA sequence (**Supplementary Table S3**). In the case of the plasmid expected to encode *SSTR1*, Sanger sequencing instead reported a sequence that is identical to *SSTR2* (**Supplementary Table S4**). This independent analysis using Sanger sequencing confirms that these two ORFs were not captured in PPLseq because they were apparently not present in the library. Thus we were able to demonstrate that PPLseq accurately describes ORF library content.

### Accuracy of PPLseq

We next analyzed the accuracy of sequences obtained using PPLseq at the single nucleotide level. We found that all but the above six absent ORFs and *PTH1R* (which is discussed separately below) had complete coverage over their entire ORFs (**Figure 3a,b**, **Supplementary Figure S4**, **Supplementary Data S4**). Variant calling demonstrated that of the 307 full length ORFs all but eleven were sequenced without deviation from the expected reference ORF sequence (i.e., with 100% sequence identity; **Figure 3c**). For these eleven genes, between one and four apparent variants were reported, and most of these 20 apparent variants were single nucleotide polymorphisms (**Figure 3d,e**, **Supplementary Data S5**). We analyzed site-level quality scores and the ratio of alternate and reference bases (expressed as the fraction of alternate base reads (AR fraction)) for the sites with reporter variants (**Figure 3f,g**, **Supplementary Data S5**). Quality scores were either very small (<1, for all variants with AR fractions <100%) or very large (>2000, for all variants with AR fractions of 100%). We first investigated the variants with high quality scores using Sanger sequencing (*BB3* (C927T), *DRD3* (A527G), and *PTH1R* (A1714G); **Figure 3h**). In the case of *BB3* and *DRD3*, we confirmed the substitutions observed in PPLseq that were not documented in the reference sequences for this library (**Supplementary Table S5, top**). The substitution in *BB3* was synonymous (F309F). The substitution in *DRD3* was non-synonymous encoding for an amino acid change in the protein relative to the deposited reference sequence (E176G; however, a glycine at this position corresponds to the canonical coding sequence). In the case of *PTH1R*, we found that the library strain encoded a canonical gene that was 22 amino acids longer than the reference sequence (**Supplementary Table S6**), also providing an explanation for the incomplete coverage noted above. We next investigated the variants with low quality scores using Sanger sequencing (**Figure 3h**). We found that none of these variants were present in the library (**Supplementary Table S5, bottom**). These findings substantiate post-calling filtering based on quality scores for reporting of final sequences filtered variants (**Figure 3i-k**). As a further test for the variant calling methodology, we introduced ten single nucleotide variants (insertions, deletions, and substitutions) into the reference ORF sequences and asked whether these could be identified in the workflow. This was indeed the case for all introduced variants with high quality scores and AR fractions (**Supplementary Figure S6**, **Supplementary Data S6,S7**). Overall, these results demonstrate single nucleotide accuracy sequencing in the efficient PPLseq method.

**Figure 3.**
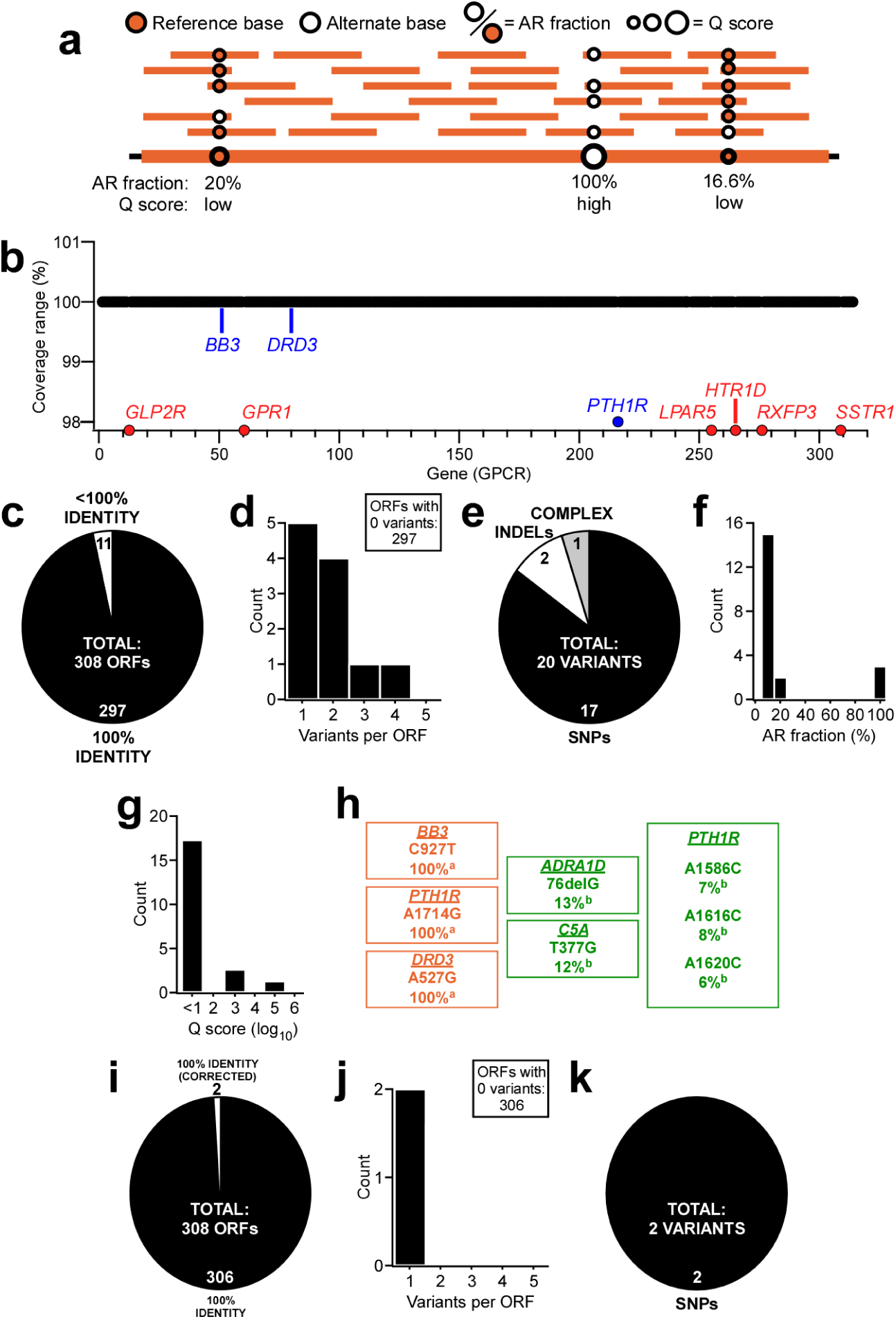
Accuracy of PPLseq. (a) Variant calling parameters (quality scores and AR fractions) in an illustrative ORF. (b) Sequence range of each GPCR ORF that is covered by mapped reads. Red spheres indicate genes from Figure 2 with limited reads and zero coverage. Blue spheres indicate receptors with variants discussed in the **Main Text** and below. (c) The number of ORFs recovered with 100% or less than 100% sequence identity (i.e., recovered with and without variants). (d,e) Distribution of variants per ORF and variant types. (f,g) Distribution of AR fractions and Q factors for all variants. (h) Variants verified using Sanger sequencing. Variants shown in orange were confirmed whereas variants shown in green were not confirmed. a: high quality score (>100); b low quality score (<1). (j) The number of ORFs recovered with 100% sequence identity after variant filtering. (j,k) Distribution of quality score filtered variants per ORF and their variant types.

### GPCR library generation supported by PPLseq

We next utilized PPLseq in the generation of a new library that contains 246 GPCRs fused to the FP mCherry (**Figure 4a**). This library is complementary to available GPCR collections that aim at understanding pharmacological and signaling properties [37–39, 42] and to the best of our knowledge the largest assembly of GPCRs with a FP fused to their C-terminus. For library generation, we applied the Type IIS restriction enzyme-based high throughput ‘Golden Gate’ gene insertion method [48]. We found that 280 of the 314 human GPCRs were amenable to this technique using two restriction enzymes selected with a hierarchical approach (SapI (229 ORFs) and BsmBI (51 ORFs), see **Materials and Methods**). We amplified each receptor in individual PCRs using ORF-specific primers from the pooled GPCR library and inserted the PCR products into a vector containing mCherry. Anticipating limitations of the cloning technique (see below), we generated six sub-libraries: three sub-libraries for each insertion reaction with each sub-library containing the complement of restriction enzyme-specific GPCR ORFs (**Figure 4a**). PPLseq then proved both efficient and powerful in understanding library generation. A single NGS reaction was sufficient to characterize each sub-library consisting of a pool of dozens of ORFs. The composition of each sub-library was resolved in detail. No reads could be mapped in any of the sub-libraries to *GLP2R*, *GPR1*, *HTR1D*, *LPAR5*, and *RXFP3* (**Supplementary Data S8-S13**), which we showed were not present in the starting library as reference sequences. Similar numbers of inserted genes were identified in the individual sub-libraries and insertion rates were generally >80% reflective of the high efficiency of the Golden Gate method (**Figure 4b,c**). As expected, distribution of genes across the groups of sub-libraries were mutually exclusive (i.e., genes inserted with SapI were absent from the BsmBI sub-libraries, and *vice versa*) (**Figure 4d**). The sequences of the inserted genes were verified through variant calling (**Figure 4e**, **Supplementary Data S14-S19**). Of a total of 679 inserted ORFs, 551 (79.5%) were identical to the library sequence after quality score post-calling variant filtering. The remainder exhibited between one and three single nucleotide variants that were predominantly single nucleotide polymorphisms (SNPs) (**Figure 4f,g**). This analysis allowed assembly of a final library from the sub-libraries for functional studies (**Supplementary Data S20**). The final library encompassed 246 mCherry-fused receptors with 100% sequence coverage and without variants, with the exception of synonymous substitutions in *GABBR1* (G930A, GABAB1-A310A), *GPR150* (T36G, GPR150-P12P), and *RXFP1* (G909A, RXFP1-G303G), as well as one non-synonymous substitution in *CHRM5* (T368C, M5R-V123A), which were included to maximize ORF numbers. Thus, PPLseq was valuable for understanding the efficiency of library construction at scale, including the specificity of amplification and insertion reactions, and compilation of a new receptor library.

**Figure 4.**
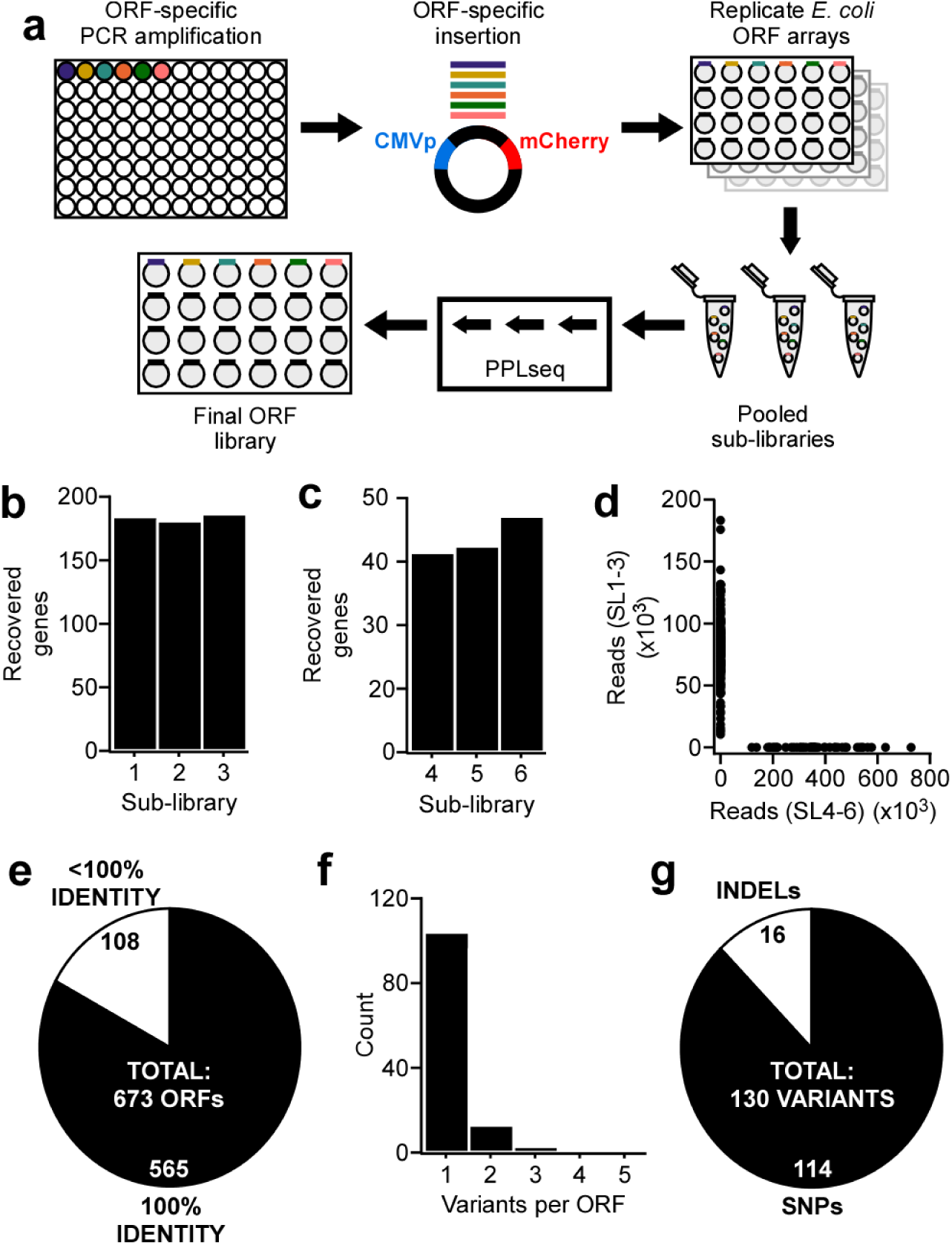
PPLseq-assisted generation of a new GPCR library. (a) Library generation through PCR reactions, Golden Gate insertion, sub-library fractionation (three sub-libraries per enzyme), and PPLseq. (b,c) Inserted GPCR ORFs recovered in six sub-libraries for insertion reactions utilising SapI (b, sub-library 1-3; out of 229 ORFs) and BsmBI (c, sub-library 4-6; out of 51 ORFs). Recovery criteria was 100% sequence coverage. (d) Mutually exclusive read distribution according to aggregated sub-libraries for the inserted 679 GPCR ORFs. (e) The number of ORFs recovered with 100% or less than 100% sequence identity (i.e., recovered with and without filtered variants). (f,g) Distribution of quality score filtered variants per ORF and their variant types. SL: sub-library.

### Expression profiles in new GPCR library

The 246 GPCR-mCherry-fusion ORFs were expressed in HEK293 cells to determine their levels using confocal microscopy and single cell analysis. Excluding controls, a total of 7.2 million single cells across 12,300 microscopy fields-of-view were analyzed corresponding to an average of 29,300 single cells per ORF (**Supplementary Data S21**). We first optimized transfection conditions using two receptors with different expression levels. In microscopy images (**Figure 5a**, **Supplementary Figure S7a,b**), we quantified single cell fluorescence intensity and transfection efficiencies (**Figure 5b**, **Supplementary Figure S7c-f**). Recognizing the limitations of transient transfections, we found that transfection with 25 ng of GPCR-mCherry-fusion vectors resulted in cell transfection rates comparable to higher doses but markedly lower aggregation (**Supplementary Figure S7c-f**). Thus, this plasmid amount was chosen for experiments. Expression was observed for most receptors. For instance, expression levels of 234 receptors exceeded a threshold of five standard deviations (SD) above negative control wells (**Figure 5c**). We found that receptor fluorescence intensities varied by nearly ten-fold between the lowest and highest expressing receptor (**Figure 5c**) and were robust between biological replicates (**Figure 5d**). The receptor expression profiles were further analyzed. First, we identified the ten most highly expressed receptors and examined their tendency to aggregate (**Figure 5d**, **Supplementary Figure S8**). We observed this for some receptors. For instance, receptors for the pituitary adenylate cyclase-activating polypeptide (*ADCYAP1R1*) or melanocyte-stimulating hormones (*MC5R*) were largely membrane localized, whereas, for instance, the EP subtype receptor for prostaglandin E₂ (*PTGER2*) was found to form aggregates (**Supplementary Figure S8**). Second, we correlated expression level profiles to ORF length and GPCR classification. Class C or B2 GPCRs generally exhibited lower expression levels, in line with their complex architectures that include large extracellular domains (**Figure 5e**). Finally, we cross-referenced expression to information about the availability of ligands and atomic resolution structures (**Figure 5f,g**). Among the receptors that express at moderate to high levels (in the 50^th^ percentile), we identified 80 receptors for which between zero and two ligands have been reported (**Figure 5f**), and 63 receptors for which between zero and two structures have been reported (**Figure 5g**). Collectively, these results demonstrate the applicability of the new GPCR library in expression level profiling towards the identification of, e.g., robustly expressed receptors that are membrane localized or understudied for further biochemical or structural investigation.

**Figure 5.**
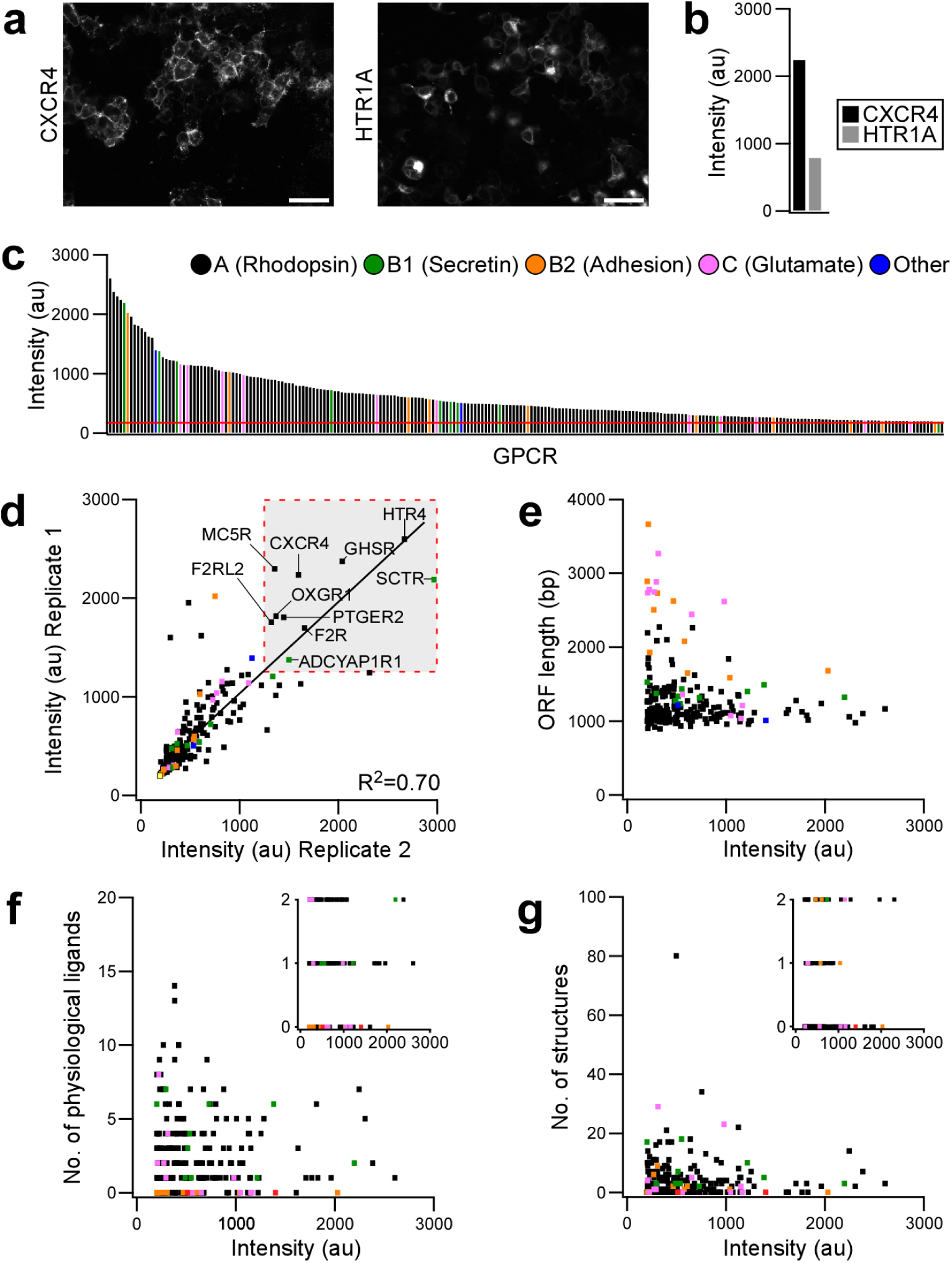
Visualisation of the GPCR ORF library. (a) Representative confocal microscopy images of HEK293 cells expressing *CXCR4-mCherry* and *HTR1A-mCherry*. Scale bars are 10 µm. (b) Average fluorescence in *CXCR4-mCherry* and *HTR1A-mCherry* transfected cells. (c) Fluorescence intensity distribution across GPCR-mCherry library (replicate 1) color-coded by GPCR family. Red line denotes the average intensity of negative controls. (d) Correlation between imaging replicates. In addition, the ten most highly expressed receptors are labelled. Receptor classes are color-coded as in (c). (e) Analysis of ORF length and fluorescence intensity. Receptor classes are color-coded as in (c). (f) Analysis of known physiological ligands and fluorescence intensity. Inset: Intensities of receptors with zero to two known ligands. (g) Analysis of available atomic resolution structures and fluorescence intensity. Inset: Intensities of receptors with zero to two available structures.

## Discussion

GOF screens utilizing ORF libraries are powerful tools for large-scale exploration of protein function. Whereas NGS is routinely applied to quantify enrichment in pooled screens, including those studying ORFs [5, 7, 25–27], few studies utilized modern sequencing methods to examine library integrity. NGS facilitated whole-genome ORF screens of model organisms [10, 17] and pathogens [49–51] as well as determined coverage in libraries of small protein domains [4, 20] and full length proteins or plasmids [13, 52]. However, in these studies, sequencing focused either on library intermediates or partial libraries and/or if sequence deviations were reported these were not resolved through independent validation. In addition, screens performed before the widespread adoption of NGS necessarily relied on libraries that were less extensively characterized. This underscores the need for a framework towards library sequencing at scale and with single-nucleotide accuracy.

We systematically established super-family library NGS by integrating sequence uniqueness mapping, pooled library preparation, and characterization of reference and new libraries. Uniqueness analysis, which has been previously applied to genomes and transcriptomes [43, 44], provided a measure to gauge fidelity of read alignments. This analysis showed that even in large super-family libraries only a limited number of genes, in particular gene isoforms, are beyond the reach of short-read NGS with ≥ 150 cycles. In PPLseq, read mapping and variant calling established on a reference library demonstrated complete recovery (i.e., identification of all ORFs in the library preparation) and high accuracy (i.e., identification of sequence variants that were then independently validated). Whereas there was general agreement between the predicted and observed mean depth, we noted variation in depth between ORFs. We attribute this to differences in input amount, e.g. due to varying plasmid copy numbers, and we can in turn conclude that the sequencing depth chosen here allowed adequate library representation. Finally, sequencing of sub-libraries using PPLseq facilitated the generation of a new GPCR library consisting of 246 mCherry-fused receptors.

We envision several potential use cases of PPLseq. In general, the method directly reports key elements of library integrity, in particular missing sequences or sequence variations. As ORF-specific barcoding is not a part of the workflow, this method is -within the limitations imposed by sequence uniqueness-directly applicable to historic libraries, including pooled libraries. With the exception of ORF libraries of the propagation host (*E. coli*), the method is agnostic to the origin of the sequence elements. PPLseq will support generation of new libraries using high-throughput gene ‘shuffling’ methods, such as the Golden Gate or Gateway techniques. Despite the robustness of these methods, it may be advisable to validate newly generated libraries. Integration of PPLseq in library generation can be somewhat more complex than characterization of existing libraries as in some instances multiple clonal replicates of the same ORF are analyzed. We showed that this can be achieved through scalable sub-libraries, such as three sub-libraries in our application of the Golden Gate method. Finally, since many gene knock-in or knock-out techniques rely on gene fragments or plasmids containing genome homologous sequences, applications of PPLseq are not limited to overexpression screens.

Assisted by PPLseq, we have generated, to the best of our knowledge, the largest GPCR-FP fusion library. Experimental characterization of GPCRs, e.g. their expression levels or sub-cellular localization, generally rely on well-by-well library arrays, despite recent successes in the development of pooled assays [53, 54]. The GPCR fusion proteins were transiently expressed in HEK293 cells and their expression was visualized using confocal microscopy. Whereas our methodology is distinct from work using surface immunostaining and FACS in the same cell type [41], we consistently observed that many receptors express at low levels and a correlation of expression and receptor classes. GPCR expression profiles may be harnessed in future studies that go beyond the development of PPLseq that was the focus of this work. These studies may focus on identification of receptors with efficient membrane localization for structural studies [40], the generation of fluorescent sensors [55, 56], or for incorporation in chimeric receptors [57–59]. We note that overexpression systems utilizing sequence-optimized genes do not recapitulate certain aspects of endogenous receptor expression, such as their dependence on DNA sequence features [41].

PPLseq is not without limitations. Knowledge of the expected sequences in the library is required for read mapping; thus, the methods is not suited for analysis of libraries with unknown content. As PPLseq relies on short-read sequencing, it is limited when resolving splice variants, but these can identified *a priori* through uniqueness analysis. Similarly, domains fused to ORF inserts are verified in vectors prior to ORF insertion. In future work, some of these limitations may be addressed through combinations with long-read sequencing methods [52, 54].

Although an ever-growing complement of structural features of proteins and their complexes can be inferred from sequences, it is to date still *in situ* experiments that allow understanding protein function, molecular interactions, and cellular localization. Elucidating this complement of properties across entire superfamilies remains challenging and requires high-throughput approaches. We present a barcode-free sequencing workflow with single nucleotide accuracy that will support plasmid library characterization in a variety of contexts. Just like NGS has revolutionized our understanding of the genome, this accurate and accessible methodology sets the stage for the use of NGS in studies that analyze coding and non-coding sequences at scale using large and flexible libraries.

## Author contributions

Conceptualization, R.T.S., H.J.; Funding Acquisition, H.J.; Methodology, R.T.S., K.H.C., J.F., S.M., G.B.G., H.J.; Project Administration, H.J.; Investigation, R.T.S., K.H.C., J.F., G.B.G., H.J.; Data Curation, R.T.S., K.H.C., G.B.G., H.J.; Supervision, H.J.; Visualization, R.T.S., K.H.C., G.B.G., H.J.; Writing - Original Draft, R.T.S., H.J.; Writing - Review and Editing, R.T.S., H.J.

## Acknowledgements

We thank Josh Dubowsky and Michael Roach for helpful comments and Matthew Popp, Josh Dubowsky, and Daniel Carrillo Baez for assistance with experiments. This study was supported by grants of the Australian Research Council (FT200100519 and DP200102093, to H.J.), the National Health and Medical Research Council (APP1187638, to H.J.). This research was supported by Flinders University through the Deep Thought High-Performance Computing environment. The Australian Regenerative Medicine Institute is supported by grants from the State Government of Victoria and the Australian Government. The EMBL Australia Partnership Laboratory (EMBL Australia) is supported by the National Collaborative Research Infrastructure Strategy (NCRIS) of the Australian Government. Imaging was performed in the CellScreen SA screening center at Flinders University.

## Materials and Methods

### Library curation

ORF sequences were compiled from public repositories in August 2021 (**Supplementary Table S1**). In the case of the GPCR library, curation was based on sequences deposited in the Addgene Inc. repository (Addgene Kit #1000000068). Of the available sequences, the third ‘depositor partial sequence’ column in the sequence dataset contained intact ORFs for all but 13 GPCRs. Sequences were considered intact if a start codon was present and if their lengths were indicative of intact codons (multiples of three nucleotides). For the remaining 13 sequences, incomplete terminal codons were manually corrected or entire sequences completed using sequencing information provided by Addgene Inc.. For the kinase and TF ORF library, sequence curation was based on sequences provided by the DNASU Plasmid Repository at the Biodesign Institute of the Arizona State University. These sequences were manually amended to correct reading frames and/or exclude stop codons (for clones in the ‘closed’ ‘fusion format’) or added terminal codons (for clones in the ‘open’ ‘fusion format’) (**Supplementary Table S1**).

### Uniqueness analysis and deduplication of ORF libraries

The kinase and TF libraries contained duplicate sequences (most commonly because in some cases the same gene was present in two library clones) that had to be removed prior to conducting a meaningful uniqueness analysis. Deduplication was performed with USEARCH (v11) using a fast variant of the UCLUST algorithm [60]. In the process of deduplication, input sequences were sorted by decreasing length and an identity threshold of 90% was applied. Cluster centroids were retained for further analysis. In the kinase and TF library, 135 and 44 sequences were removed, resp.. Unique ORF sequences are provided in **Supplementary Data S1-S3**.

To test for sequence uniqueness, ORF sequences were then concatenated into long strings separated with non-nucleotide characters. Strings were then searched 5’ to 3’ in forward direction and backward direction (in instances where forward searching did not report a match) using sliding windows of pre-defined length (50, 75, 150, or 300 b). Occurrence of the window sequence in another location of the string was registered in binary format. The number and length of non-unique sequence segments along with the corresponding ORF names was determined using a level crossing algorithm. Analysis was performed using macros written in C in Igor Pro (v6.22).

For sequence pairs reported to contain non-unique segments, sequence overlap (using pairwise alignments), naming and presence of splice variants (using nucleotide BLAST searches and GeneCards) and internal repeats (using Tandem Repeats Finder v4.09 with default parameters [61] were manually verified.

### Regrowth of GPCR collection

The PRESTO-Tango GPCR collection was a kind gift from Bryan Roth and colleagues [39]. *E. coli* glycerol stocks were obtained from Addgene.org (Addgene Kit #1000000068) and replica-plated using sterile disposable 96-pin replicator plates (Ciro Manufacturing Corp.) onto single-well plates (242811, Thermo Fisher Scientific Inc.) containing LB-ampicillin agar. The plates were incubated overnight at 37°C and replica-plated the following day for imaging on a white light imager (Gel Logic 212 Pro, Carestream Health Inc.). Inoculation spots for each plate were then scraped in 5 ml of LB-ampicillin medium into sterile tubes for downstream plasmid purification using four mini-scale silica columns. Each plate resulted in an elution volume of >150 μl and DNA concentrations between 135 and 360 ng/μl.

### Validation of genes with incomplete coverage (Sanger sequencing)

The library clones that should encode *LPAR5* and *SSTR1* yielded growing cultures (**Supplementary Figure S3**) but either zero or only minimal read numbers limited to short sequence stretches could be mapped to these sequences (**Supplementary Figure S4**, **Supplementary Data S4**). To investigate this, clones were restreaked from the original GPCR library and a clonal culture was grown overnight at 37°C in 2 ml LB-ampicillin medium. Purified plasmids were analyzed using Sanger Sequencing (AGRF Ltd.) using forward and reverse primers (primers 1 and 2, **Supplementary Table S7**).

### Validation of genes with low reads and incomplete coverage (PCR)

The library clones that should encode *GLP2R*, *GPR1*, *HTR1D*, and *RXFP3* did not yield growing cultures (**Supplementary Figure S3**) and no reads could be mapped to these sequences (**Supplementary Figure S4**, **Supplementary Data S4**). We thus utilized PCR with gene-specific forward and reverse primers to test for their presence in the library pool (primers 12 to 19, **Supplementary Table S7**; *TSHR* served as a control, primers 20 to 21, **Supplementary Table S7**). Reactions contained 2.5 μl 10x Pfu polymerase buffer (B600003, Sangon Biotech (Shanghai) Co. Ltd.), template DNA (containing ∼30 pg target gene if present), 1 μl forward primer (10 μM stock concentration), 1 μl reverse primer (10 μM stock concentration), 0.5 μl dNTP mix (10 mM stock concentration; U151B, Promega Corp.), 0.5 μl Pfu polymerase (5 u) (B600003, Sangon Biotech (Shanghai) Co. Ltd.) in a total volume of 25 µl. PCR cycling parameters were 1. 2 minutes at 95°C, 2. 30 seconds at 95°C, 3. 30 seconds at 60°C, 4. 1 minute at 72°C, 5. Repeat steps 2 to 4 a further 23 times. Reactions were performed in 200 μl PCR tubes in a T100 thermal cycler (Bio-Rad Laboratories Inc.).

### NGS sample preparation

NGS samples were prepared in low DNA binding tubes (0030108418, Eppendorf SE). Typically, >2 μg of purified plasmids were collected in a final volume of 30 μl. These pools were created by combining material isolated either from the four replica plates (original ORF collection) or from twelve individual cloning plates (plasmids created with the Golden Gate method). In the case of cloning plates, each sub-library pool contained up to one clone per ORF (see below). The amount of input material from each plate was chosen such that the number of clones on each plate were represented with equal amounts in the pool, except for sequencing of the initial GPCR library, where read numbers for ORFs with >2 reads on a partial plate were normalized in comparison and visualization. Libraries were prepared using TruSeq DNA Nano kit (20015964, Illumina Inc.) with 100 ng starting material and 350 bp fragment size. Libraries were sequenced on the NovaSeq6000 platform (Illumina Inc.) with a paired-end read length of 150 b. Sequencing was provided by JS LINK Inc.

### NGS analysis

Tabulated ORF sequences including short extensions into the plasmid backbone (33 b each at the 5’ and 3’ end) served as reference sequences for NGS analysis. The reference sequence collection also included a plasmid backbone sequence for the mapping of non-ORF reads. Between 23 and 44 million filtered reads were processed for each of the NGS samples, and between 87 and 96% of reads were mapped to the ORF libraries (including plasmid backbones). Paired-end sequencing reads were filtered and aligned to reference sequences with Snippy (v4.6.0) [62] and BWA (v0.7.17-r1188) [63] to generate an alignment output file. Alignment statistics, including mapped read numbers, coverage, and sequencing depth, were extracted from the alignment output file using SAMtools (v1.18) [64]. Variant calling was performed with FreeBayes (v1.3.6) [65] with default parameters (minimum mapping quality ≥60, base quality ≥13, coverage ≥4, ALT-supporting reads ≥2, ALT allele fraction ≥5%) to identify single or multi-nucleotide polymorphisms and indels. Raw variant data from FreeBayes without further Snippy filtering were extracted using BCFtools (v1.19) [64]. Variant quality scores correspond to Phred-scaled probabilities that a polymorphism exists at this site. Final data were compiled in a spreadsheet where each gene was annotated with read numbers, depth and coverage. Variants were then reviewed to identify true deviations from the references sequences defined as those with quality scores above the default Snippy threshold of 100 (see **Main Text** for details). Not specified are variants in the plasmid backbone.

### Variant validation (Sanger sequencing)

Selected sequence variants identified by PPLseq were verified using Sanger sequencing also to support the quality score-based post-calling filtering (**Supplementary Table S5**). Clones were prepared directly from the original GPCR library and a clonal culture was grown overnight at 37°C in 2 ml LB-ampicillin medium. Purified plasmids were analyzed using Sanger Sequencing (AGRF Ltd.) using primers 3 to 7 (**Supplementary Table S7**).

### Vectors for Golden Gate method

Golden Gate insertions were performed int modified pcDNA 3.1(−) vector (V79520, Thermo Fisher Scientific Inc.). In this vector, six recognition sites for three Type IIS restriction enzymes (two sites for BsaI, one for BbsI, and three for SapI) were previously removed using site-directed mutagenesis [42]. In addition, the large section of the vector comprising the F1 origin of replication and aminoglycoside resistance cassette was removed to reduce vector size. Next, two receiver cassettes were designed and synthesized that contain (i) the FP mVenus [66] optimized for bacterial expression and under the control of a bacterial promoter (J23115, https://parts.igem.org/Part:BBa_J23115), (ii) SapI or BsmBI restriction sites flanking mVenus, and (iii) the FP mCherry positioned for C-terminal fusion to the inserted GPCR. mVenus served as a removable marker protein to identify colonies in which the Golden Gate reaction had proceeded. The cassettes were obtained as GBlock gene fragments (Integrated DNA Technologies Inc.) and inserted into the new vector *via* restriction enzyme cloning. The cassettes were first amplified by PCR using Pfu polymerase (B600003, Sangon Biotech (Shanghai) Co. Ltd.). The vector and cassettes were then digested with XbaI and HindIII restriction enzymes (R0145 and R0104, New England Biolabs Inc.) and ligated with T4 DNA Ligase (M180B, Promega Corp.). Sequences were verified using Sanger sequencing (AGRF Ltd.).

### Primers and PCRs for Golden Gate insertion

A script was employed for automated primer design and screening of internal Type IIS restriction enzyme recognition sites in GPCR ORF sequences. ORF sequences were searched for recognition sites of the restriction enzymes SapI, BsmBI, BsaI, BbsI, BseRI, and PaqCI. The prevalence of recognition sites across the entire library was considered to determine an enzyme hierarchy so that the maximal number of sequences could be cloned with the smallest number of enzymes. This hierarchy was SapI > BsmBI, BsaI > BbsI > BseRI > PaqCI. Given the small number of GPCRs incompatible with either SapI or BsmBI (34 GPCRs), sequences within that group were excluded from the workflow. Primers were automatically designed with appropriate annealing sites specific to the start and end of each ORF with a melting temperature of >64°C. The primers contained flanking overhangs with the respective Type IIS sites for Golden Gate insertion and were synthesized in 96-well plates on the scale of 500 pmoles (Integrated DNA Technologies Inc.).

The pooled GPCR library was used as the template for gene amplification. Reactions contained 2.5 μl 10x Pfu polymerase buffer (B600003, Sangon Biotech (Shanghai) Co. Ltd.), 1 μl pooled template DNA (10 ng), 1 μl forward primer (10 μM stock concentration), 1 μl reverse primer (10 μM stock concentration), 0.5 μl dNTP mix (10 mM stock concentration, U151B, Promega Corp.) and 0.5 μl Pfu polymerase (5 u) (B600003, Sangon Biotech (Shanghai) Co. Ltd.) in a total volume of 25 μl. PCR cycling parameters were 1. 2 minutes at 95°C, 2. 30 seconds at 95°C, 3. 30 seconds at 55°C, 4. 5 minutes at 72°C, 5. Repeat steps 2 to 4 a further 29 times, and 6. 10 minutes at 72°C. PCRs were performed in 96-well PCR plates (HSS9601, Bio-Rad Laboratories Inc.) in a T100 thermal cycler (Bio-Rad Laboratories Inc.)

### Golden Gate method

Golden Gate reactions utilized SapI or BsmBI restriction enzymes (R0739 and R0569, New England Biolabs Inc.) depending on ORF enzyme compatibility. Reactions contained 2 μl T4 ligase buffer (C126A, Promega Corp.), 1 μl receiver plasmid (100 ng), 2 μl unpurified PCR product, 0.5 μl (5 u) restriction enzyme, and 1 μl (3 u) T4 Ligase (M180B, Promega Corp.) in a total volume of 20 μl. Reactions were performed in 96-well PCR plates (3441-00, Scientific Specialists Inc.). Reactions were incubated at room temperature (RT) for 30 minutes (SapI) or thermocycled (BsmBI) with the following parameters: 1. 2 minutes at 37°C, 2. 3 minutes at 16°C, 3. Repeat steps 1 to 2 a further 11 times, 4. 5 minutes at 50°C, and 5. 10 minutes at 80°C in the T100 thermal cycler. In the next step, 0.2 μl of each reaction was then transformed into 20 μl NEB Turbo competent cells (C2984H, New England Biolabs Inc.) for 20 minutes on ice and plated on large-scale well plates containing LB-ampicilin to incubate overnight at 37°C.

For each transformed Golden Gate reaction, three colonies were manually picked and grown individually in 200 μl LB-ampicllin media overnight at 37°C. In the next step, 50 μl of each overnight culture were obtained from the first colony of each Golden Gate reaction, pooled, and purified using standard silica columns for NGS. This was repeated for second and third colonies. At the same time and for each individual culture, 50 μl of each overnight culture was mixed with 25 μl of sterile glycerol (50%) and stored at −80°C in 96-well plates.

### Assembly of sequenced GPCR fusion protein library

Successful insertion of the intact GPCR ORF into the plasmid was verified by PPLseq and defined as 100% sequence coverage with no variants (see **Main Text** for exceptions). Sequence-verified clones were grown starting from the glycerol stock. If more than one colony was sequence-verified, colonies with the highest number of reads or colonies from the sub-library with maximum validated ORFs were selected. For each colony, the glycerol stock was tip touched into 200 μl of LB-ampicllin media and incubated overnight at 37°C with shaking at 600 rpm. 50 μl from each well was mixed with 25 μl of sterile glycerol (50%) and stored at −80°C.

The final glycerol stock was then transferred into 96-well deep well plates containing 1.5 ml LB-ampicllin media using the sterile disposable 96-pin replicator plates. The cultures were sealed with a breathable foil (BF-400-S, Axygen Inc.) and incubated overnight at 37°C with shaking at 600 rpm. Plasmids were purified using 96-well plasmid purification kits (B519196, Sangon Biotech (Shanghai) Co. Ltd.). The method followed the manufacturer’s recommendations with minor modification: centrifugation steps were performed at 3,350 x g, bacteria were pelleted for 3 minutes, solutions were mixed by pipetting, and lysates were clarified for 40 minutes.

### Mammalian cell culture and transfection

HEK293 cells were cultured in Dulbecco’s Modified Eagle’s Medium (DMEM) (1195073, Thermo Fisher Scientific Inc.) supplemented with 10% fetal bovine serum (FBS) (26140079, Thermo Fisher Scientific Inc.), 100 U ml^−1^ penicillin and 0.1 mg ml^−1^ streptomycin (15140122, Thermo Fisher Scientific Inc.) in a humidified incubator in 5% CO_2_ atmosphere at 37°C. Reverse transfections utilizing polyethyleneimine (PEI) were performed as described previously [67, 68] in 96-well PhenoPlates (6055302, Revvity Inc.) coated with poly-*L*-lysine (P8920, Sigma-Aldrich Corp.). For each receptor, 250 ng of DNA (in the library screen: 90% of plasmid without an inserted ORF and 10% ORF plasmid) was diluted to a total volume of 25 µl with pre-warmed Opti-MEM (OM) (31985070, Thermo Fisher Scientific Inc.). Additionally, for each receptor, 1 µg polyethyleneimine (PEI) was diluted to a total volume of 25 µl with prewarmed OM. The combined DNA-OM and PEI-OM solutions were incubated for 20 minutes at RT. The mixture was then added to 2 × 10^4^ cells in 100 µl antibiotic-free DMEM containing 5% FBS. Six hours post incubation, the original medium was replaced with 150 µl complete DMEM.

### Staining and imaging

Cells were washed 48 hours after transfection with 100 μl prewarmed phosphate buffered saline (PBS) (20012050, Thermo Fisher Scientific Inc.). Cells were then incubated with 50 μl of 1x working concentration of Cell Navigator Cell Plasma Membrane Stain (AAT-22682, AAT BioQuest Inc) and 16 μM Hoechst 33342 (14533, Sigma-Aldrich Corp.) for 15 minutes at 37°C. Cells were then washed a further two times with 100 μl prewarmed PBS and fixed in 50 μl 3% paraformaldehyde (C0042, ProSciTech Pty Ltd.) for 20 minutes at RT. After fixation, cells were washed one more time and maintained in PBS at 4°C until imaging. Imaging was performed on an Operetta automated confocal microscope with Harmony (v4.1) image acquisition and image analysis software (Revvity Inc.). Confocal fluorescence images were obtained using excitation/emission wavelengths of 475 ± 15 / 525 ± 25 nm (Cell Navigator), 380 ± 20 / 445 ± 35 (Hoechst), and 570 ± 10 / 615 ± 25 (mCherry). In this analysis, 25 fields of view were recorded per well at a magnification of 20x with the confocal plane located through the cell midline. Cells transfected with plasmids lacking ORFs or left untransfected served as negative controls.

### Image analysis

Raw images were imported into a dedicated local network environment and image analysis was performed in the Columbus (v2.5) image data storage and analysis system on an Omero (v4.0) and Acapella (v3.2) server (all from Revvity Inc.). In total, 16,100 images and 9.8 million individual cells were analyzed from 644 individual wells (25 fields of view per well). Of these, 2,400 images and 2.1 million cells were from positive and negative control wells, 12,300 images and 7.2 million single cells were from the GPCR-ORF library, and the remainder from further calibration and reference wells. The nucleus of individual cells was identified using nuclear counter staining Hoechst. Cells on the edges of the images were discarded. Following nuclei localization, cell bodies were segmented using the plasma membrane stain with the cytoplasm identification method “C” in Columbus. mCherry fluorescence intensity was measured in the whole cell as a normalized value to the cell area and this value was used to classify cells as transfected or not transfected. Cells with intensity values greater than the mean + 5 SD of the negative control wells (on a per plate basis) were considered transfected. Only wells with >100 transfected cells were included in the analysis (this applied to 238 ORFs in either of the biological replicates).

### GPCR ligand and structure analysis

Receptor names were standardized by mapping HGNC symbols to corresponding IUPHAR identifiers using the Guide to Pharmacology (IUPHAR) database (https://www.guidetopharmacology.org/GRAC/GPCRListForward?class=A). Formatting artifacts, such as HTML tags and symbols were removed prior to downstream analysis. For each receptor, the following was retrieved from GPCRdb (human receptors only): receptor family classification, the number of available structures (https://gpcrdb.org/structure/), and the number of deposited physiological ligands (https://gpcrdb.org/ligand/physiological_ligands/). Duplicate receptor-ligand entries were removed. In cases where automated look-up failed (e.g., due to formatting or nomenclature differences), information was manually retrieved from GPCRdb.

