## Supplementary Figures and Tables for "Gene library deep sequencing for protein super-family profiling"

### Supplementary Information

**Supplementary Information:**

Supplementary Tables S1-S7

Supplementary Figures S1-S8

Supplementary Data S1-S22

### Supplementary Tables

**Supplementary Table S1. ORF library sequences analyzed for uniqueness.**

| Protein Family | Total ORFs | Unique ORFs[a] | Total bases of unique ORFs | Source of sequence information |
|---|---|---|---|---|
| Human GPCRs | 314 | 314 | 422,430[b] | Addgene Inc., Kit #1000000068 |
| Human kinase | 630 | 495 | 878,907[c] | DNASU Plasmid Repository, The Biodesign Institute/Arizona State University[d] |
| Human TFs | 2037 | 1993 | 3,182,480[e] | DNASU Plasmid Repository, The Biodesign Institute/Arizona State University[f] |

[a] See **Materials and Methods** for deduplication methods

[b] Excluding added epitopes, cleavage site and transactivator domain

[c] Excluding stop codons (for clones in the 'closed' 'fusion format') or added terminal codons (for clones in the 'open' 'fusion format')

[d] Harvard Institute of Proteomics (HIP) at Harvard Medical School human kinase collection

[e] Excluding stop codon for one clone in the 'closed' fusion format

[f] Human ORFeome v2 TF subcollection

**Supplementary Table S2. ORF pairs identified in uniqueness analysis.** Only pairs identified at windows lengths of 150 and 300 b are specified here.

| First ORF | Second ORF | Reasons for non-uniqueness |
|---|---|---|
| Human GPCRs | | |
| No non-unique sequences | | |
| Human kinases | | |
| *MAP2K2* | *MAP2K2* | Splice isoforms[a] |
| *MST4* | *MST (RP6-213H19.1)* | Splice isoforms[a] |
| Human TFs | | |
| *ETV3L* | *ETV3* | Paralogs[b] |
| *IFI16* | *IFI16* | Internal repeats[b] |
| *PEG3* | *ZIM2* | Shared exons[b] |
| *STAT5A* | *STAT5B* | Paralogs[a] |
| *ZKSCAN3* | *ZKSCAN4* | Paralogs[b] |
| *ZNF354A* | *ZNF354B* | Paralogs[b] |
| *ZNF552* | *ZNF814* | Paralogs[b] |
| *ZNF814* | *ZNF814* | Internal repeats[b] |
| *ZNF765* | *ZNF813* | Paralogs[b] |
| *ZSCAN5A* | *ZSCAN5B* | Paralogs[b] |

[a] Identified at 150 and 300 b window lengths

[b] Identified at 150 b window length

**Supplementary Table S3. Sanger sequencing results for *LPAR5*.** The determined and deposited proteins sequences are 100% identical but encoded using different codon usages. Plasmid was sequenced with primers 1 and 2 (**Supplementary Table S7**).

| | |
|---|---|
| Assembled codon-altered nucleotide sequence | GGGGTGGTGCGTTAACTTAAGCTTGGTACCGAGCTCGGATCCACT<br>AGTCCAGTGTGGTGGAATTCTGCAGATATCCAGCACAGTGGCGGC<br>CGCGCCACCATGAAGACGATCATCGCCCTGAGCTACATCTTCTGC<br>CTGGTATTCGCCGACTACAAGGACGATGATGACGCCAGCATCGATA<br>TGCTCGCTAATTCTAGTAGTACTAATTCATCCGTTCTGCCGTGCCCC<br>GACTACAGGCCGACCCACCGCCTGCACTTGGTGGTATATTCCTTG<br>GTGCTCGCCGCCGGGCTTCCTCTCAATGCCTTGGCTCTGTGGGT<br>GTTCCTGAGGGCCCTCCGAGTGCACAGCGTTGTGAGTGTATACAT<br>GTGCAACCTTGCAGCCAGCGATCTGCTGTTCACCTTGTCACTTCC<br>CGTGAGGCTTAGTTACTACGCCCTTCATCACTGGCCCTTTCCCGAC<br>CTCCTCTGTCAGACTACCGGAGCCATCTTCCAGATGAATATGTACG<br>GATCATGCATTTTCCTGATGCTCATTAATGTGGACCGCTACGCAGC<br>AATCGTTCATCCCCTCAGATTGAGACATCTTAGACGACCACGGGTG<br>GCTCGGCTCCTGTGCCTCGGTGTGTGGGCTCTGATCTTGGTATTT<br>GCTGTTCCCGCGGCAAGGGTGCACCGGCCCTCACGATGCCGGTA<br>TCGCGATTTGGAAGTTAGACTCTGCTTCGAATCATTTTCTGACGAA<br>CTGTGGAAGGGAAGGCTTTTGCCCCTTGTGTTGCTCGCTGAGGC<br>CTTGGGGTTTCTCTTGCCCCTTGCAGCGGTGGTCTATAGTTCAGG<br>CAGAGTTTTCTGGACCCTGGCAAGGCCCGACGCAACACAGTCCC<br>AAAGAAGGCGCAAAACAGTGCGCTTGTTGTTGGCCAACCTGGTCA<br>TCTTTCTCCTGTGTTTTGTCCCCTATAATAGTACTCTGGCAGTGTAC<br>GGGCTCCTGAGGAGCAAACTTGTGGCCGCCTCTGTGCCAGCTAG<br>GGATCGAGTGAGAGGAGTTCTCATGGTCATGGTGCTTTTGGCAGG<br>CGCTAATTGCGTCCTGGACCCTCTTGTGTATTACTTTTCCGCCGAA<br>GGTTTTCGCAACACACTGAGGGGGTTGGGCACTCCGCATAGGGC<br>CAGGACGAGTGCAACTAATGGGACACGGGCTGCACTCGCGCAGA<br>GTGAAAGATCTGCTGTCACCACCGATGCCACAAGACCCGACGCTG<br>CATCCCAGGGACTCCTCCGGCCCAGCGACAGTCACAGTCTGTCTA<br>GCTTCACACAGTGCCCTCAGGACAGCGCATTGATCGATACCGGTG<br>GACGCACCCCACCCAGCCTGGGTCCCCAAGATGAGTCCTGCACC<br>ACCGCCAGCTCCTCCCTGGCCAAGGACACTTCATCGACCGGTGA<br>GAACCTGTACTTCCAGCTAAGATTAGATAAAAGTAAAGTGATTAACA<br>GCGCATTAGAGCTGCTTAATGAGTCGATNAAGCC |
| Determined protein sequence (GPCR only, equiv. to NP_001136433) | MLANSSSTNSSVLPCPDYRPTHRLHLVVYSLVLAAGLPLNALALWVFL<br>RALRVHSVVSVYMCNLAASDLLFTLSLPVRLSYYALHHWPFPDLLCQ<br>TTGAIFQMNMYGSCIFLMLINVDRYAAIVHPLRLRHLRRPRVARLLCLG<br>VWALILVFAVPAARVHRPSRCRYRDLEVRLCFESFSDELWKGRLLPLV<br>LLAEALGFLLPLAAVVYSSGRVFWTLARPDATQSQRRRKTVRLLLANL<br>VIFLLCFVPYNSTLAVYGLLRSKLVAASVPARDRVRGVLMVMVLLAGA<br>NCVLDPLVYYFSAEGFRNTLRGLGTPHRARTSATNGTRAALAQSERS<br>AVTTDATRPDAASQGLLRPSDSHSLSSFTQCPQDSAL |
| Deposited protein sequence (Addgene #66421, equiv. to NP_001136433) | MLANSSSTNSSVLPCPDYRPTHRLHLVVYSLVLAAGLPLNALALWVFL<br>RALRVHSVVSVYMCNLAASDLLFTLSLPVRLSYYALHHWPFPDLLCQ<br>TTGAIFQMNMYGSCIFLMLINVDRYAAIVHPLRLRHLRRPRVARLLCLG<br>VWALILVFAVPAARVHRPSRCRYRDLEVRLCFESFSDELWKGRLLPLV<br>LLAEALGFLLPLAAVVYSSGRVFWTLARPDATQSQRRRKTVRLLLANL<br>VIFLLCFVPYNSTLAVYGLLRSKLVAASVPARDRVRGVLMVMVLLAGA<br>NCVLDPLVYYFSAEGFRNTLRGLGTPHRARTSATNGTRAALAQSERS<br>AVTTDATRPDAASQGLLRPSDSHSLSSFTQCPQDSAL |

**Supplementary Table S4. Sanger sequencing results for *SSTR1*.** The determined and deposited proteins sequences are non-identical as described in the **Main Text**. Plasmid was sequenced with primers 1 and 2 (**Supplementary Table S7**).

| | |
|---|---|
| Assembled nucleotide sequence | GGGGTGGNGCGTTAACTTAAGCTTGGTACCGAGCTCGGATCCACT<br>AGTCCAGTGTGGTGGAATTCTGCAGATATCCAGCACAGTGGCGGC<br>CGCGCCACCATGAAGACGATCATCGCCCTGAGCTACATCTTCTGC<br>CTGGTATTCGCCGACTACAAGGACGATGATGACGCCAGCATCGATA<br>TGGACATGGCCGATGAGCCTCTGAACGGTTCCCACACCTGGCTGA<br>GCATACCTTTCGATTTGAATGGTTCAGTCGTGTCAACGAACACGAG<br>CAATCAGACCGAACCATATTATGACCTGACTAGCAATGCCGTTTTGA<br>CTTTCATTTACTTTGTGGTGTGCATCATCGGATTGTGCGGAAATACC<br>CTCGTCATTTACGTCATCCTGCGGTATGCAAAGATGAAAACTATTAC<br>CAACATTTACATCCTCAATCTGGCCATCGCGGATGAACTGTTTATGC<br>TGGGACTGCCTTTCCTGGCGATGCAGGTGGCACTCGTTCACTGG<br>CCTTTCGGAAAAGCTATATGTAGGGTCGTAATGACTGTAGACGGAA<br>TTAACCAGTTCACTTCAATTTTCTGTTTGACTGTTATGAGTATTGAC<br>CGCTATCTTGCCGTCGTCCACCCGATAAAGTCAGCCAAGTGGCGG<br>AGACCAAGAACCGCTAAAATGATCACAATGGCTGTGTGGGGGGTT<br>AGCCTGCTGGTTATCCTGCCAATTATGATCTATGCAGGCCTTAGGT<br>CAAATCAATGGGGCCGATCTAGTTGCACAATCAATTGGCCCGGCG<br>AGAGTGGAGCATGGTACACGGGGTTCATTATCTATACATTCATTCTC<br>GGCTTCCTGGTACCACTCACAATTATATGTCTGTGTTACCTGTTTAT<br>AATTATTAAGGTAAAAAGCAGCGGGATACGGGTGGGAAGTTCAAAA<br>CGAAAGAAGTCTGAAAAAAAAGTGACCAGAATGGTGTCTATCGTG<br>GTGGCTGTATTTATTTTTTGTTGGTTGCCGTTCTACATTTTCAATGTC<br>AGTTCAGTTTCTATGGCCATAAGCCCAACCCCTGCACTTAAAGGTA<br>TGTTTGATTTCGTGGTCGTTCTGACCTACGCCAACAGTTGCGCCAA<br>TCCCATTCTGTACGCCTTTCTTAGCGACAACTTCAAGAAAAGCTTT<br>CAAAACGTTTTGTGCCTGGTTAAGGTGAGTGGAACTGATGACGGC<br>GAGAGATCCGACAGTAAGCAGGACAAAAGTAGACTTAATGAGACC<br>ACCGAGACCCAACGGACCCTCCTCAATGGGGACCTGCAGACCTC<br>CATCATCGATACCGGTGGACGCACCCCACCCAGCCTGGGTCCCCA<br>AGATGAGTCCTGCACCACCGCCAGCTCCTCCCTGGCCAAGGACA<br>CTTCATCGACCGGTGAGAACCTGTACTTCCAGCTAAGATTAGATAA<br>AAGTAAAGTGATTAACAGCGCATTAGAGCTGCTTAATGAGTTCGAT<br>GAANCCC |
| Determined protein sequence (GPCR only, equiv. to NP_001041) | MDMADEPLNGSHTWLSIPFDLNGSVVSTNTSNQTEPYYDLTSNAVLT<br>FIYFVVCIIGLCGNTLVIYVILRYAKMKTITNIYILNLAIADELFMLGLPFLA<br>MQVALVHWPFGKAICRVVMTVDGINQFTSIFCLTVMSIDRYLAVVHPIK<br>SAKWRRPRTAKMITMAVWGVSLLVILPIMIYAGLRSNQWGRSSCTINW<br>PGESGAWYTGFIIYTFILGFLVPLTIICLCYLFIIIKVKSSGIRVGSSKRKK<br>SEKKVTRMVSIVVAVFIFCWLPFYIFNVSSVSMAISPTPALKGMFDFVV<br>VLTYANSCANPILYAFLSDNFKKSFQNVLCLVKVSGTDDGERSDSKQD<br>KSRLNETTETQRTLLNGDLQTSI |
| Deposited protein sequence (Addgene #66502, equiv. to NP_001040) | MFPNGTASSPSSSPSPSPGSCGEGGGSRGPGAGAADGMEEPGRN<br>ASQNGTLSEGQGSAILISFIYSVVCLVGLCGNSMVIYVILRYAKMKTAT<br>NIYILNLAIADELLMLSVPFLVTSTLLRHWPFGALLCRLVLSVDAVNMFT<br>SIYCLTVLSVDRYVAVVHPIKAARYRRPTVAKVVNLGVWVLSLLVILPIV<br>VFSRTAANSDGTVACNMLMPEPAQRWLVGFVLYTFLMGFLLPVGAIC<br>LCYVLIIAKMRMVALKAGWQQRKRSERKITLMVMMVVMVFVICWMPF<br>YVVQLVNVFAEQDDATVSQLSVILGYANSCANPILYGFLSDNFKRSFQ<br>RILCLSWMDNAAEEPVDYYATALKSRAYSVEDFQPENLESGGVFRNG<br>TCTSRITTL |

**Supplementary Table S5. Sanger sequencing validation.** Comparison of Sanger sequencing results and deposited sequences. Plasmids were sequenced with primers 3 to 7 (**Supplementary Table S7**). Green highlights indicate variants discussed in the main text.

| Gene | Ref / bp / Alt | |
|---|---|---|
| **Above threshold variants** | | |
| *BB3* | C927T | |
| sequencing | | TACCATTCATTCACTTCTCAAACCTATGTAGACCCCTCTGCCATGCATTT**T**ATTTTCACCATTTTCTCTCGGGTTTTGGCTTTCAGCAATTCTTGCGTAAA |
| deposited as Addgene #66228 | | TACCATTCATTCACTTCTCAAACCTATGTAGACCCCTCTGCCATGCATTT**C**ATTTTCACCATTTTCTCTCGGGTTTTGGCTTTCAGCAATTCTTGCGTAAA |
| *DRD3* | A527G | |
| sequencing | | ACTGGCCTTTGCTGTGTCCTGCCCTCTTCTGTTTGGCTTTAATACCACAG**G**GGACCCCACTGTCTGCTCCATCTCCAACCCTGATTTTGTCATCTACTCTT |
| deposited as Addgene #66270 | | ACTGGCCTTTGCTGTGTCCTGCCCTCTTCTGTTTGGCTTTAATACCACAG**A**GGACCCCACTGTCTGCTCCATCTCCAACCCTGATTTTGTCATCTACTCTT |
| **Sub-threshold variants** | | |
| *C5A* | T377G | |
| sequencing | | TAGTATCCTGCCCTCTCTGATTCTGCTCAACATGTACGCGTCAATACTCC**T**TCTCGCAACCATTAGCGCGGACAGGTTTCTCTTGGTGTTCAAGCCCATTT |
| deposited as Addgene #66232 | | TAGTATCCTGCCCTCTCTGATTCTGCTCAACATGTACGCGTCAATACTCC**T**TCTCGCAACCATTAGCGCGGACAGGTTTCTCTTGGTGTTCAAGCCCATTT |
| *ADRA1D* | G76- | |
| sequencing | | TCTCCTTTGAGGGGCCCCGACCAGACAGCAGCGCCGGAGGAAGCAGTGCT**G**GGGGGGGGGGTGGGTCCGCCGGAGGCGCTGCTCCCTCCGAGGGGCCGGCG |
| deposited as Addgene #66215 | | TCTCCTTTGAGGGGCCCCGACCAGACAGCAGCGCCGGAGGAAGCAGTGCT**G**GGGGGGGGGGTGGGTCCGCCGGAGGCGCTGCTCCCTCCGAGGGGCCGGCG |
| *PTH1R* | A1586C | |
| sequencing | | CGGACTTGGACTGCCTTTGAGCCCTAGACTTCTGCCGACTGCCACCACTA**A**CGGTCACCCCCAACTGCCGGGCCACGCCAAGCCAGGGACACCGGCTCTGG |
| deposited as Addgene #66489 | | CGGACTTGGACTGCCTTTGAGCCCTAGACTTCTGCCGACTGCCACCACTA**A**CGGTCACCCCCAACTGCCGGGCCACGCCAAGCCAGGGACACCGGCTCTGG |
| *PTH1R* | A1616C | |
| sequencing | | TCTGCCGACTGCCACCACTAACGGTCACCCCCAACTGCCGGGCCACGCCA**A**GCCAGGGACACCGGCTCTGGAAACCCTGGAAACGACTCCTCCCGCAATGG |
| deposited as Addgene #66489 | | TCTGCCGACTGCCACCACTAACGGTCACCCCCAACTGCCGGGCCACGCCA**A**GCCAGGGACACCGGCTCTGGAAACCCTGGAAACGACTCCTCCCGCAATGG |
| *PTH1R* | A1620C | |
| sequencing | | CCGACTGCCACCACTAACGGTCACCCCCAACTGCCGGGCCACGCCAAGCC**A**GGGACACCGGCTCTGGAAACCCTGGAAACGACTCCTCCCGCAATGGCGGC |
| deposited as Addgene #66489 | | CCGACTGCCACCACTAACGGTCACCCCCAACTGCCGGGCCACGCCAAGCC**A**GGGACACCGGCTCTGGAAACCCTGGAAACGACTCCTCCCGCAATGGCGGC |

**Supplementary Table S6. Sequence analysis of *PTH1R*.** Plasmid was sequenced with primer 7 (**Supplementary Table S7**).

| | |
|---|---|
| Raw Sanger sequencing result (3’ end of *PTH1R*) | GGGGNCNGCCTATTGAAATGCTTTTCAACTCCTTCCAGGGATTTTTCG<br>TGGCCATCATCTATTGTTTTTGCAACGGCGAGGTTCAGGCAGAGATTA<br>AAAAATCTTGGTCCAGATGGACCCTTGCGCTGGACTTCAAGAGAAAG<br>GCGCGATCTGGGAGCAGTTCCTATTCTTACGGACCCATGGTGAGTCA<br>CACCTCCGTCACCAACGTTGGCCCCCGGGTCGGACTTGGACTGCCT<br>TTGAGCCCTAGACTTCTGCCGACTGCCACCACTAACGGTCACCCCCA<br>ACTGCCGGGCCACGCCAAGCCAGGGACACCGGCTCTGGAAACCCT<br>GGAAACGACTCCTCCCGCAATGGCGGCACCCAAAGACGACGGGTTC<br>CTGAATGGGTCATGCTCCGGTTTGGATGAGGAAGCCTCTGGACCGGA<br>GAGACCCCCCGCTCTGCTGCAAGAAGAATGGGAAACCGTGATGATC<br>GATACCGGTGGACGCACCCCACCCAGCCTGGGTCCCCAAGATGAGT<br>CCTGCACCACCGCCAGCTCCTCCCTGGCCAAGGACACTTCATCGAC<br>CGGTGAGAACCTGTACTTCCAGCTAAGATTAGATAAAAGTAAAGTGATT<br>AACAGCGCATTAGAGCTGCTTAATGAGGTCGGAATCGAAGGTTTAACA<br>ACCCGTAAACTCGCCCAGAAGCTAGGTGTAGAGCAGCCTACATTGTAT<br>TGGCATGTAAAAAATAAGCGGGCTTTGCTCGACGCCTTAGCCATTGAG<br>ATGTTAGATAGGCACCATACTCACTTTTGCCCTTTAGAAGGGGAAAGC<br>TGGCAAGATTTTTTACGTAATAACGCTAAAAGTTTTAGATGTGCTTTACT<br>AAGTCATCGCGATGGAGCAAAAGTACATTTAGGTACACGGCCTACAGA<br>AAAACAGTATGAAACTCTCGAAAATCAATTAGCCTTTTTATGCCAACAA<br>GGTTTTTCACTAGAGAATGCATTATATGCACTCAGCGCTGTGGGGCAT<br>TTTACTTTAGGTTGCGTATTGGAAGATCAAGAGCATCAAGTCGCTAAA<br>GAAGAAAGGGAAACACCTACTACTGATAGTATGCCGCCATTATTACGA<br>CAAGCTATCGAATTATTTGATCACCAAGGTGCAGAGCCAGCCTTCTTAT<br>TCGGCCTTGAATTGATCATATGCGGATTAGAAAACAACTTAAATGGGAA<br>AGTGGGTCCGCGTACAGCCGCGCCCGTACAAAAAAAATTACGGGTCT<br>ACCATCGAGGGCCGGCCGATTCCCGGAAGANGACCCCCCCAAAAAG<br>GGGGGTTGGGGGTCCCGCCNNTTCTTTTCCCCCGGG |
| Translated region | …EMLFNSFQGFFVAIIYCFCNGEVQAEIKKSWSRWTLALDFKRKARSGS<br>SSYSYGPMVSHTSVTNVGPRVGLGLPLSPRLLPTATTNGHPQLPGHAKP<br>GTPALETLETTPPAMAAPKDDGFLNGSCSGLDEEASGPERPPALLQEEW<br>ETVM |
| Determined protein sequence (GPCR only, 593 aa, equiv. to NP_000307) | MGTARIAPGLALLLCCPVLSSAYALVDADDVMTKEEQIFLLHRAQAQCEK<br>RLKEVLQRPASIMESDKGWTSASTSGKPRKDKASGKLYPESEEDKEAPT<br>GSRYRGRPCLPEWDHILCWPLGAPGEVVAVPCPDYIYDFNHKGHAYRR<br>CDRNGSWELVPGHNRTWANYSECVKFLTNETREREVFDRLGMIYTVGY<br>SVSLASLTVAVLILAYFRRLHCTRNYIHMHLFLSFMLRAVSIFVKDAVLYSG<br>ATLDEAERLTEEELRAIAQAPPPPATAAAGYAGCRVAVTFFLYFLATNYYW<br>ILVEGLYLHSLIFMAFFSEKKYLWGFTVFGWGLPAVFVAVWVSVRATLANT<br>GCWDLSSGNKKWIIQVPILASIVLNFILFINIVRVLATKLRETNAGRCDTRQ<br>QYRKLLKSTLVLMPLFGVHYIVFMATPYTEVSGTLWQVQMHYEMLFNSF<br>QGFFVAIIYCFCNGEVQAEIKKSWSRWTLALDFKRKARSGSSSYSYGPM<br>VSHTSVTNVGPRVGLGLPLSPRLLPTATTNGHPQLPGHAKPGTPALETLE<br>TTPPAMAAPKDDGFLNGSCSGLDEEASGPERPPALLQEEWETVM |
| Deposited protein sequence (571 aa, Addgene #66489) | MGTARIAPGLALLLCCPVLSSAYALVDADDVMTKEEQIFLLHRAQAQCEK<br>RLKEVLQRPASIMESDKGWTSASTSGKPRKDKASGKLYPESEEDKEAPT<br>GSRYRGRPCLPEWDHILCWPLGAPGEVVAVPCPDYIYDFNHKGHAYRR<br>CDRNGSWELVPGHNRTWANYSECVKFLTNETREREVFDRLGMIYTVGY<br>SVSLASLTVAVLILAYFRRLHCTRNYIHMHLFLSFMLRAVSIFVKDAVLYSG<br>ATLDEAERLTEEELRAIAQAPPPPATAAAGYAGCRVAVTFFLYFLATNYYW<br>ILVEGLYLHSLIFMAFFSEKKYLWGFTVFGWGLPAVFVAVWVSVRATLANT<br>GCWDLSSGNKKWIIQVPILASIVLNFILFINIVRVLATKLRETNAGRCDTRQ |

| | |
|---|---|
| | QYRKLLKSTLVLMPLFGVHYIVFMATPYTEVSGTLWQVQMHYEMLFNSF<br>QGFFVAIIYCFCNGEVQAEIKKSWSRWTLALDFKRKARSGSSSYSYGPM<br>VSHTSVTNVGPRVGLGLPLSPRLLPTATTNGHPQLPGHAKPGTPALETLE<br>TTPPAMAAPKDDGFLNGSCSGL |

**Supplementary Table S7. Sequencing and amplification primers used in this study.**

| Primer number, name and direction | Sequence | Purpose |
|---|---|---|
| 1 T7, forward | TAATACGACTCACTATAGGG | Sequencing |
| 2 *tTA*, reverse | CTTCTGGGCGAGTTTACGGG | Sequencing |
| 3 *C5A*, forward | TCCCGATATCTTGGCATTGGTC | Sequencing |
| 4 *BB3*, forward | ACCCTTTACAAAAGCACCCTGAA | Sequencing |
| 5 *DRD3*, forward | GGTGGCCACCTTGGTGATG | Sequencing |
| 6 *ADRA1D*, forward | ATGACTTTTAGGGACTTGCTTAGC | Sequencing |
| 7 *PTH1R*, forward | CAGAGGTCTCTGGAACATTGTG | Sequencing |
| 8 *LPAR5*, forward | ATGCTGGCCAATTCCAGCTCTA | Amplification |
| 9 *LPAR5*, reverse | GTCTCACTTCCAGGTCCCG | Amplification |
| 10 *SSTR1*, forward | ATGTTTCCCAACGGCACGGC | Amplification |
| 11 *SSTR1*, reverse | TACTTTGGCCACTGTAGGCC | Amplification |
| 12 *GLP2R*, forward | ATGAAACTCGGGTCTTCCAGG | Amplification |
| 13 *GLP2R*, reverse | CAGCAGGGCATAGCGATCG | Amplification |
| 14 *GPR1*, forward | ATGGAAGACTTGGAGGAAACACT | Amplification |
| 15 *GPR1*, reverse | TTGAATTCCACGGTGTCGCG | Amplification |
| 16 *HTR1D*, forward | ATGAGTCCTCTTAACCAGAGTGC | Amplification |
| 17 *HTR1D*, reverse | TGCCTTAGCCTGCCTCCAG | Amplification |
| 18 *RXFP3*, forward | ATGCAGATGGCCGACGCC | Amplification |
| 19 *RXFP3*, reverse | GTCAGAAAGAACACTGACGCG | Amplification |
| 20 *TSHR*, forward | ATGCGCCCTGCCGATCTG | Amplification |
| 21 *TSHR*, reverse | CCTGTACGCTGGTGAACCC | Amplification |

### Supplementary Figures

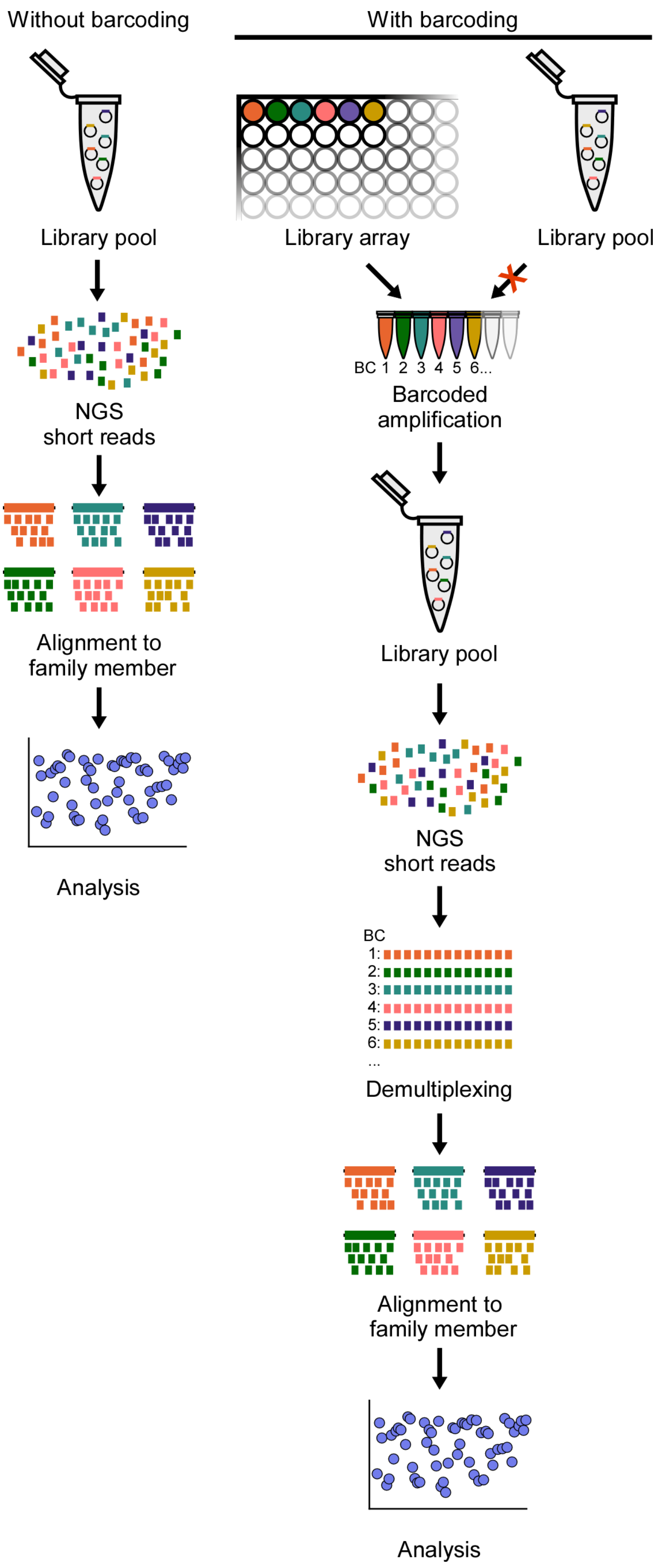


**Supplementary Figure S1. Comparison of workflow in PPLseq (left) and workflow with ORF barcoding (right).** Barcoding introduces additional steps and is incompatible with sequencing of existing libraries that are only available in pooled formats.

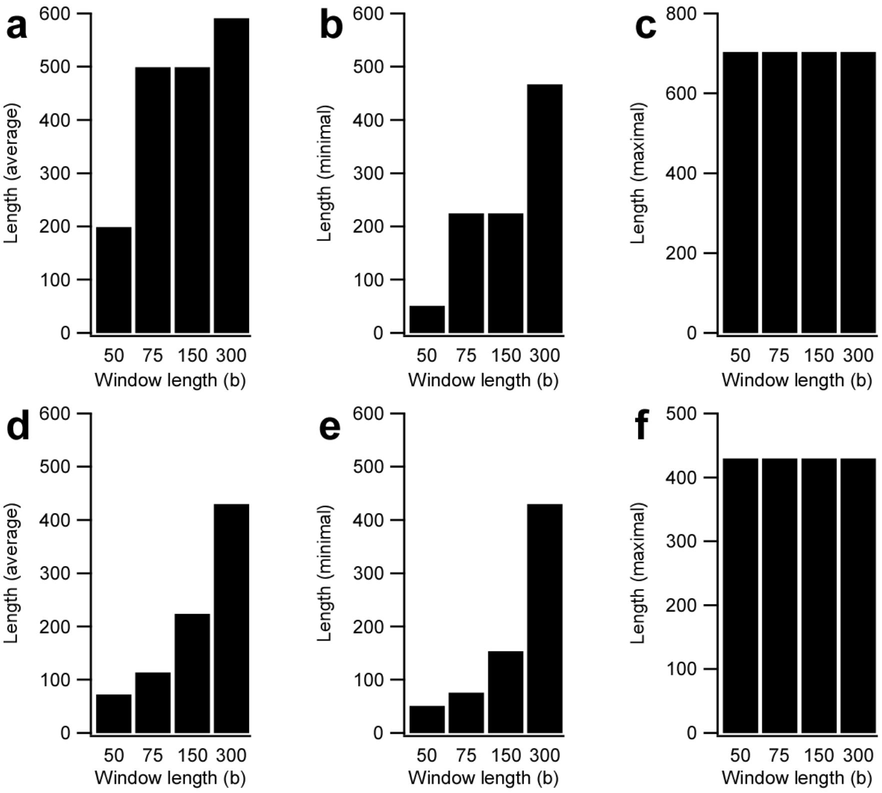


**Supplementary Figure S2. Analysis of non-unique sequence segments.** Lengths of non-unique segments in human kinases (a-c) and human TFs (d-f) expressed as averages (a, d), minimal lengths (b, e), and maximal lengths (c, f).

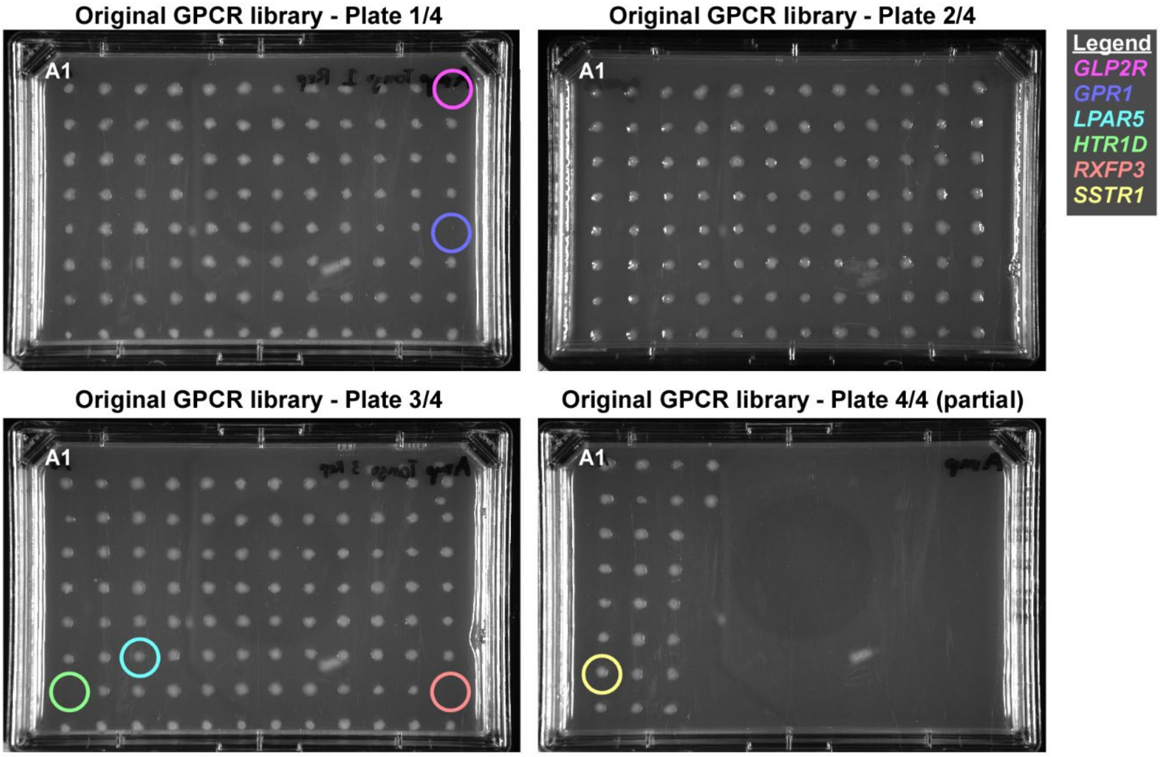


**Supplementary Figure S3. Growth cultures of replica-plated GPCR library.** ORFs not identified in PPLseq are highlighted in color.

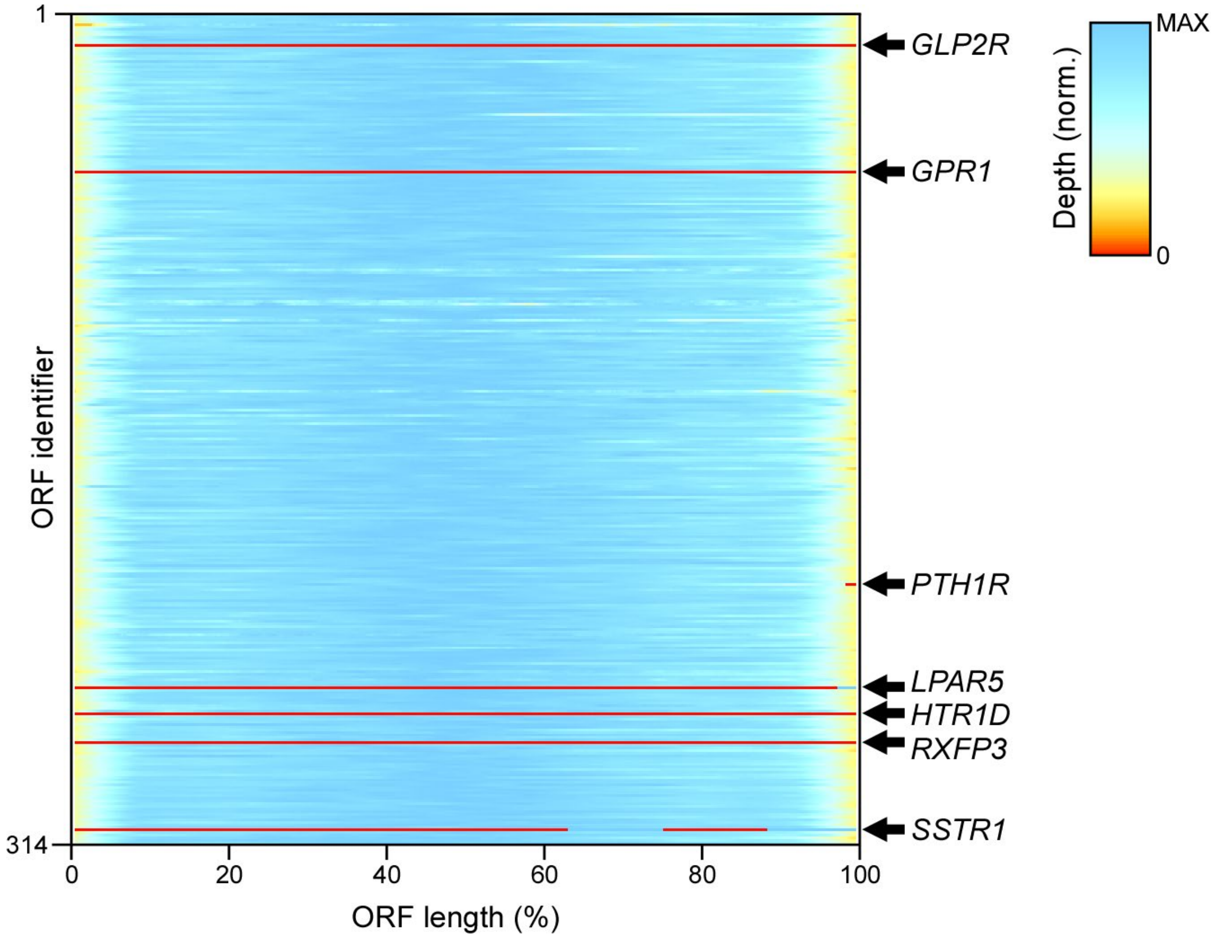


**Supplementary Figure S4. Heatmap showing sequencing depth across GPCRs.** Color scale is normalized for each receptor. Regions lacking sequence coverage are highlighted in red, with the corresponding ORFs specified on the right.

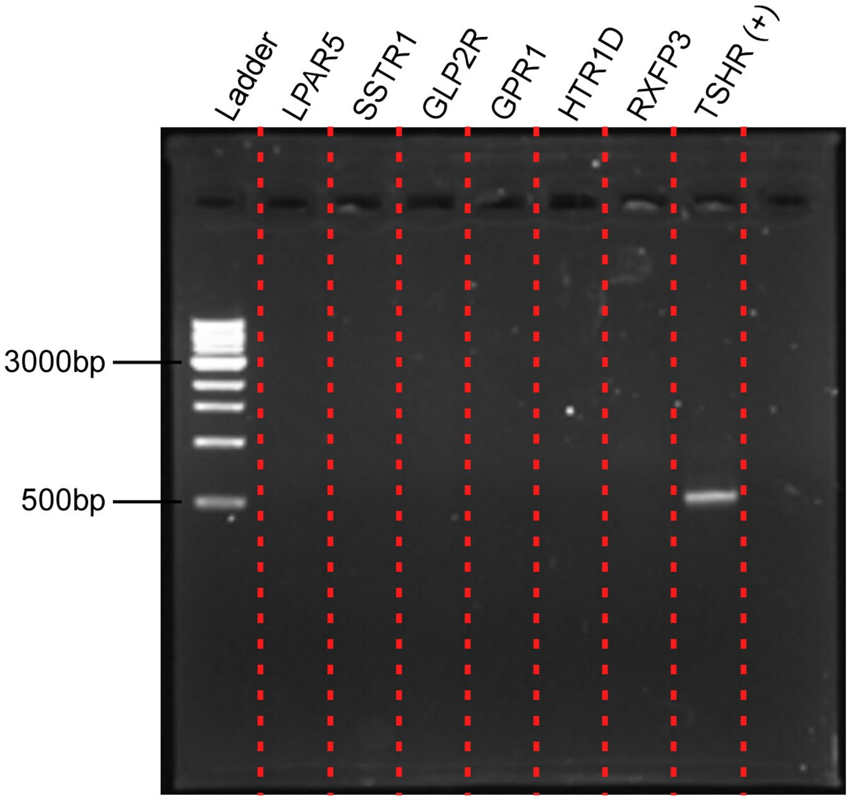


**Supplementary Figure S5. Analytical PCR reactions to test for the presence of ORFs in the GPCR library.** ORF-specific specific forward and reverse primers (**Supplementary Table S7**) were designed to amplify PCR products of ~500 bp length from *LPAR5*, *SSTR1*, *GLP2R*, *GPR1*, *HTR1D*, and *RXFP3*. *TSHR* served as the positive control.

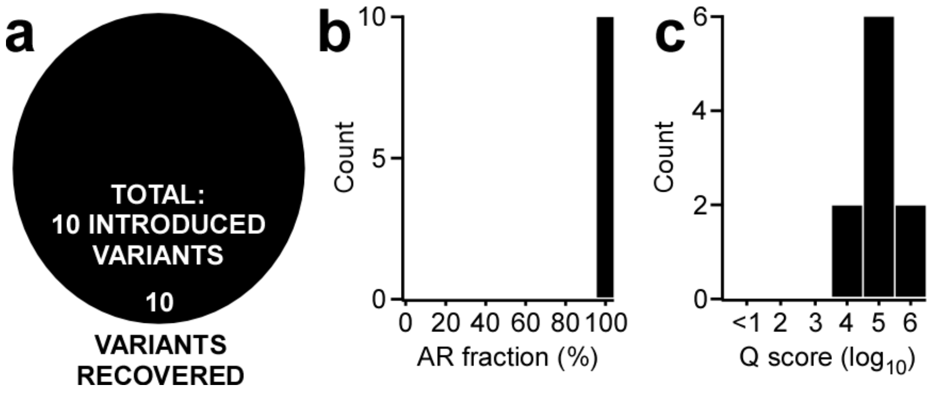


**Supplementary Figure S6. Analysis of introduced reference sequence variants.** (a) All introduce variants have been recovered. (b,c) Distribution of AR fractions and quality scores for introduced variants.

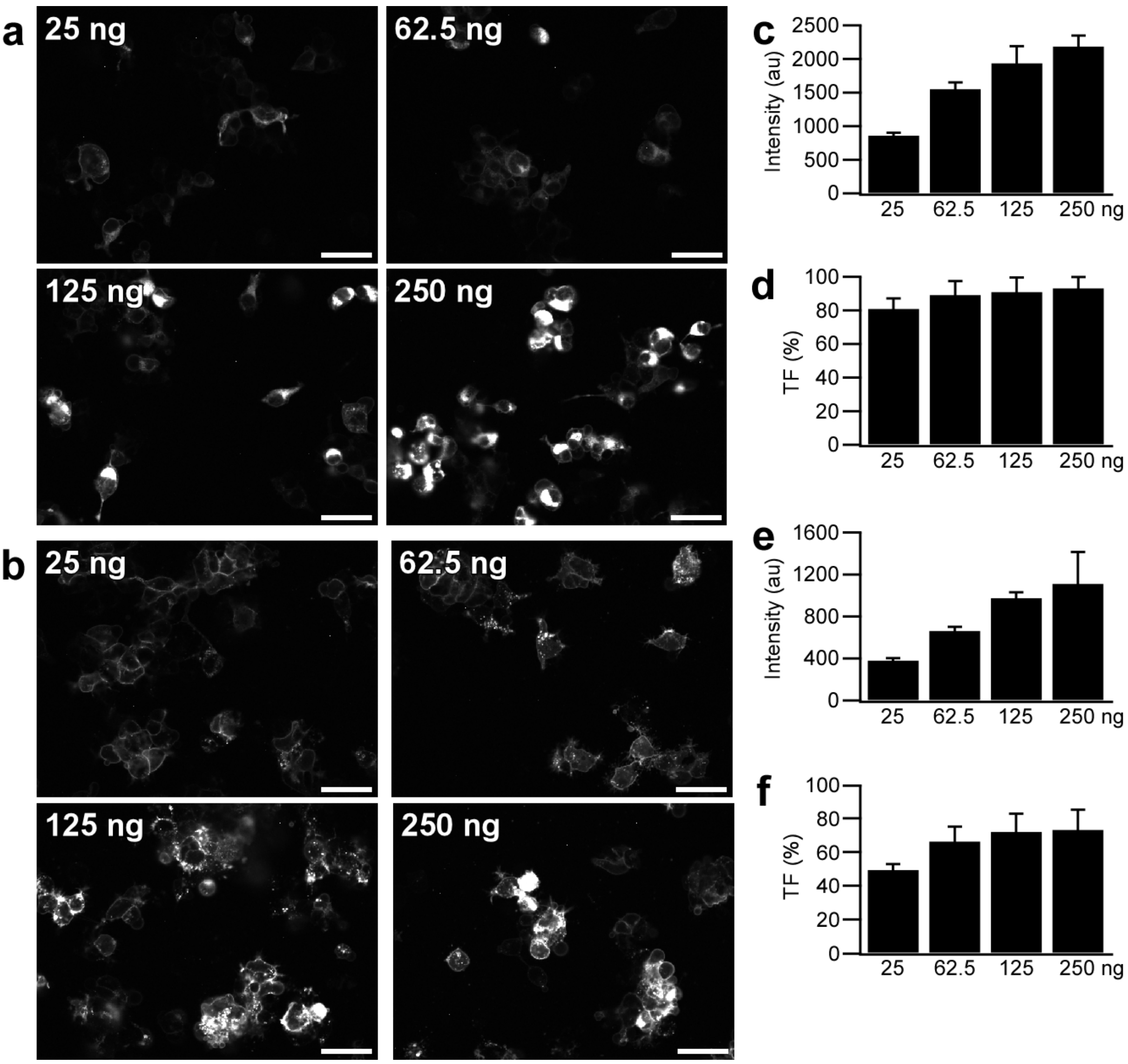


**Supplementary Figure S7. Dose-dependent expression levels of GPCRs.** (a,b) Confocal microscopy images of HEK293 cells transfected with *CXCR4-mCherry* (a) and *HTR1A-mCherry* (b) at increasing plasmid DNA amounts (25, 62.5, 125, and 250 ng). Scale bars are 10 µm. (c,e) Whole cell fluorescence intensity of transfected cells for *CXCR4-mCherry* (c) and *HTR1A-mCherry* (e) across the different dosages. Mean intensity values ± SD (n=4 wells each with 100 fields of view). (d,f) Transfection efficiency of *CXCR4-mCherry* (d) and *HTR1A-mCherry* (f) across the different dosages. TF (%): Percentage of transfected cells.

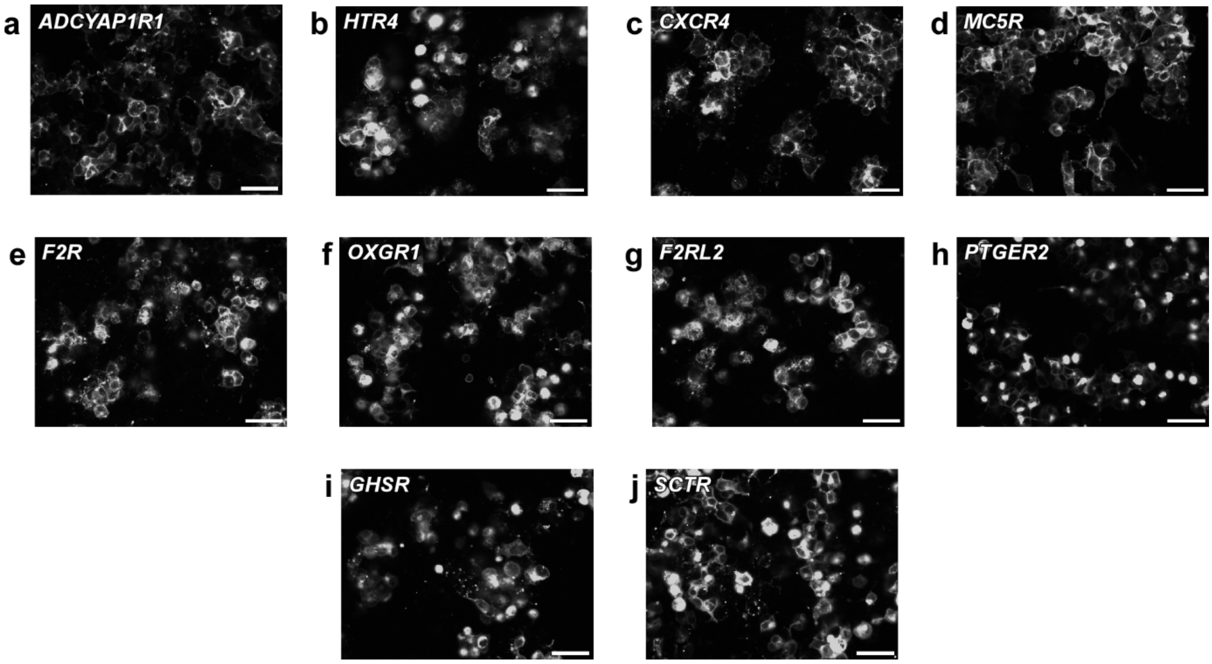


**Supplementary Figure S8. Representative confocal microscopy images for the ten highest expressed GPCRs.** Scale bars are 10 µm.
